# Gene-scale DNA and chromatin ensembles with atomic resolution

**DOI:** 10.64898/2026.11.16.766978

**Authors:** Johannes Betz, Sergio Cruz-León, Tobias Hüfner, Johannes Dopfer, Gerhard Hummer

## Abstract

The spatial organization of cellular DNA extends across a wide range of scales, from sequence-dependent base pairing to the packaging of meter-long cellular DNA into micron-size nuclei. Integrating the information from disparate experiments, molecular simulations, and theoretical models into a coherent picture of DNA and chromatin organization across scales remains challenging. Here, we introduce hierarchical chain growth for DNA (HCG-DNA) to build atomically detailed ensembles of double-stranded DNA and chromatin. Starting from complete tetranucleotide fragment libraries encoding local sequence-dependent conformations, we assemble structures of DNA up to the megabase scale at quasilinear computational cost. From HCG-DNA, the large-scale polymer elastic properties emerge, in particular the persistence length. HCG-DNA recovers electron spin resonance, atomic force microscopy, and single-molecule force spectroscopy measurements of DNA structure across scales. HCG-DNA includes nucleosomes to generate 3D atomistic chromatin ensembles constrained by cryo-electron tomography or genomic nucleosome maps. With HCG-DNA, we show how small changes in nucleosome linker length alter nucleosome packing and chromatin compaction. HCG-DNA connects atomic-level DNA and nucleosome structure to chromosome-scale spatial organization, while providing scalable ensembles for coarse-grained model development and AI training.

## Introduction

With 6 × 10^9^ base pairs (bp) in its 23 chromosome pairs, a diploid human cell packages ≈2 m of double-stranded DNA (dsDNA) into a nucleus only ≈10 µm across. The physical organization of the genome thus spans more than four orders of magnitude, from Ångström-scale molecular conformations to micrometer-scale chromatin architecture [1, 2]. Beyond encoding the genetic information, the DNA sequence governs the local structure and mechanical deformability and the interactions with ions [3–10]. These sequence-dependent properties contribute to nucleosome positioning, transcription factor binding, and polymer mechanics, linking local DNA physics to chromatin organization and biological function [11–14]. Yet how molecular DNA physics propagates across scales to generate collective mechanics and chromatin architecture remains unresolved [15–17].

Different experiments are combined to probe DNA organization across scales. Genome-wide assays such as Hi-C, MNase-seq, and ATAC-seq map chromosome contacts, nucleosome positions, and accessibility, but do not directly resolve molecular structure [18–21]. Fluorescence imaging and chromatin tracing reveal spatial organization in single cells [22–24], while in situ cryo-electron tomography (cryo-ET) localizes nucleosomes and higher-order chromatin features, yet the conformation of the DNA connecting these structures remains largely inaccessible [25, 26]. At smaller scales, crystallography, nuclear magnetic resonance (NMR), pulsed electron-electron double resonance (PELDOR; also known as double electron-electron resonance, DEER), atomic force microscopy (AFM) and single-molecule manipulation define DNA structure, conformational distributions and mechanics at near-atomic resolution [27–32].

Molecular simulations span a similar hierarchy of resolution. Atomistic molecular dynamics (MD) directly samples sequence-dependent DNA conformations and molecular interactions, and systematic simulation efforts now push to characterize larger parts of DNA sequence space [5, 7, 8, 33]. Computational cost, however, limits routine atomistic sampling to comparatively short DNA molecules [3, 6, 10, 33–35]. Coarse-grained, rigid-base and polymer models complement atomistic MD by representing nucleic acid molecular behavior through effective interactions and elastic descriptions, enabling simulations at kilobase and chromatin scales [36–44]. These approaches have provided powerful descriptions of DNA mechanics and chromatin organization [16, 39, 43, 45, 46]. However, they each make different trade-offs between molecular detail, interaction complexity and the spatial and temporal scales that can be sampled.

Hierarchical chain growth (HCG) provides a statistical-mechanical framework to build molecular conformational ensembles across scales [47]. Originally developed for intrinsically disordered proteins [48] and more recently extended to single-stranded RNA [49], HCG constructs long-chain ensembles from distributions of shorter molecular fragments while retaining the fragments’ atomistic representation [47]. The hierarchical assembly preserves detailed balance, which guarantees that arbitrary choices such as the order of the assembly do not affect the final ensemble [47, 48].

Here, we develop hierarchical chain growth for DNA (HCG-DNA) to build atomistic DNA conformations from local sequence contexts to megabase DNA and chromatin. We show that local atomistic DNA information can be propagated across orders of magnitude in length while retaining experimentally meaningful molecular and polymer properties. For HCG-DNA, we use independently generated all-atom MD libraries spanning all unique tetranucleotide contexts as molecular building blocks. We show that HCG-DNA preserves sequence-dependent conformational statistics from long atomistic simulations and recovers independent PELDOR distance distributions, AFM polymer dimensions and single-molecule DNA mechanics without fitting polymer-scale elastic parameters. HCG-DNA scales to atomically resolved 1-Mbp DNA. Combined with cryo-ET and genomic nucleosome maps, we use HCG-DNA to produce atomistic chromatin ensembles, revealing that single-base-pair changes in linker length generate periodic changes in nucleosome packing and compaction. HCG-DNA thus provides a quantitative bridge from molecular DNA physics to genome-scale organization and a scalable source of atomistic ensembles for multiscale and machine learning models.

## Materials and methods

HCG-DNA generates sequence-specific atomistic ensembles by assembling conformations sampled from molecular dynamics libraries covering all unique tetranucleotide sequence contexts. In the following, we describe in detail the generation of the fragment library, the fragment assembly algorithm and the memory-efficient implementation of HCG-DNA that allow for atomic representations at genome scale. We then show details for the comparison of HCG-DNA to diverse experiments, and the modeling of chromatin structures with HCG.

### Fragment library generation

We performed MD simulations of all 136 unique dsDNA tetramers embedded in a decamer structure. Simulations were run in Gromacs2022.4 [50] using the parmbsc1 force field [51]. We used Mamatkulov-Schwierz parameters for monovalent ions [52], which have been extensively tested on nucleic acids simulations [9, 10, 26, 34, 35, 41, 53]. The TIP3P-F water model was used to speed up conformational sampling [54]. For simulation the DNA tetramer sequence was embedded in a decamer sequence (5^′^-GCGX_1_X_2_X_3_X_4_CGC-3^′^). The starting structures were generated from HCG from a complete tetramer library generated from a previously reported MD trajectory [34] as part of initial testing. Structures were solvated in a rhombic dodecahedron box with an edge length of 8 nm containing an aqueous solution of 150 mM KCl. After steepest descent energy minimization to a maximum force of 1000 kJ mol^−1^ nm^−1^ and at most 10,000 steps, we equilibrated the system in the NVT ensemble for 500 ps with a 1 fs time step using Berendsen temperature coupling [55] to maintain 300 K. We then equilibrated in the NPT ensemble for 1 ns with a 2 fs time step using Berendsen pressure coupling to maintain 1 bar. Both equilibration steps were performed with position restraints on the DNA heavy atoms using a force constant of 1000 kJ mol^−1^ nm^−2^. We used short range real-space cut-offs of 1.2 nm for Coulombic and van-der-Waals interactions. Long-range electrostatic interactions were treated with the particle mesh Ewald scheme. For production, we ran 1.01 *µ*s simulations with a 2 fs time step using V-rescale temperature coupling [56] and C-rescale pressure coupling [57]. We saved frames every 100 ps into the fragment library. The first 10 nanoseconds of each production run were discarded from the final fragment library.

### HCG-DNA: fragment assembly algorithm

Beginning with the MD fragment libraries, we map the target DNA sequence to the corresponding overlapping tetranucleotide ensembles and randomly draw conformations from the respective libraries (Fig. 1 and Supporting Figure 1). Adjacent fragments share two base pairs (Fig. 1 left) and are assembled pairwise in a hierarchical manner. In this way, the central two base pairs of each tetramer fragment are retained in the first assembly step. Accepted conformations form the ensemble from which structures are drawn at the next level, and this procedure is repeated until the full-length sequence is assembled. During stitching, we first align two fragments on the first of the two overlapping base pairs, specifically the heavy atoms in the nucleobase and the ribose carbon atoms. If the alignment root-mean-squared deviation (RMSD) is below 0.6 Å, we check for clashes between adjacent residues using a heavy atom distance cut-off of 2 Å. If no clashes are detected in this step, we then check that the phosphorus atoms in the assembled structure are *>* 4 Å apart (Supporting Fig. 1).

**Figure 1:**
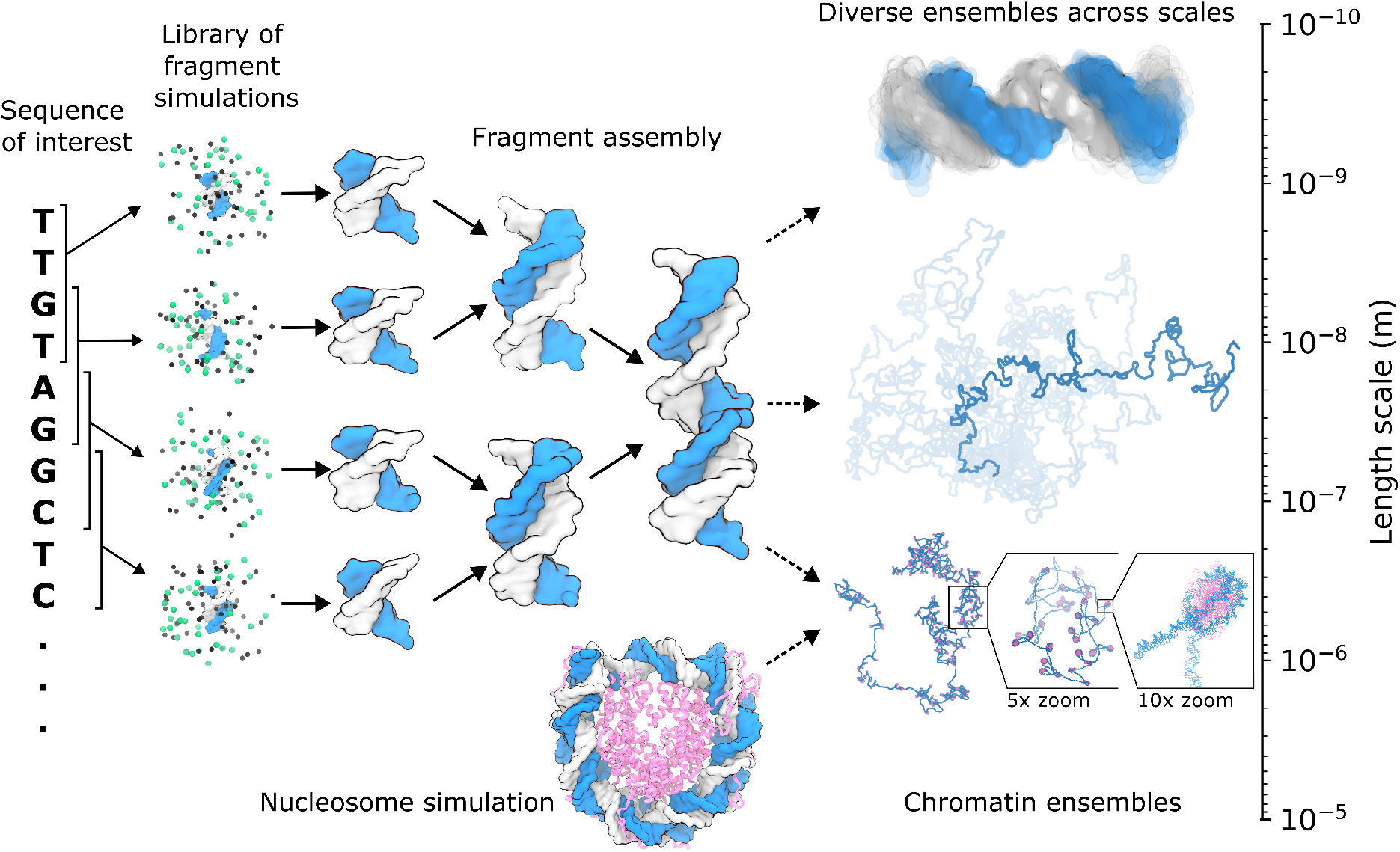
HCG-DNA hierarchically propagates atomistic DNA conformations across scales. For a sequence of interest, HCG-DNA maps tetranucleotide contexts from conformational libraries generated by all-atom molecular dynamics simulations (left). Conformations sampled from these libraries are recursively aligned and assembled through overlapping coordinates, generating increasingly long, sequence-specific DNA molecules while retaining atomic resolution (middle). Repeated sampling produces diverse structural ensembles spanning molecular to micrometer length scales (right). The same hierarchical framework incorporates nucleosome conformations sampled from atomistic simulations or experiments with HCG-generated linker DNA to construct atomically resolved chromatin ensembles (bottom). *Alt text:* Schematics and molecular visualizations illustrating the HCG-DNA process. On the left is a target sequence TTGTAGGCTC with brackets over each tetranucleotide overlapping by 2 bases. Each tetranucleotide is shown in an MD simulation system, then get combined in a pairwise manner, yielding an increasingly long chain. On the right are overlayed ensembles of a short DNA chain and a long one placed approximately on the length scale. On the bottom is a visualization of a nucleosome structure, which gets combined with DNA chains to form chromatin structures. One chromatin structure is shown with a series of zoom-ins showing the atomic detail.

In the HCG algorithm [48], the alignment and steric criteria define a binary acceptance function. Hierarchical assembly samples the product of the input fragment distributions conditioned on these constraints, avoiding the directional chain-growth bias introduced by sequential assembly. No additional energetic reweighting is applied during HCG-DNA assembly.

Since the fragment stitching does not explicitly optimize for the phosphate group of the backbone-phosphate geometry, elongated O3^′^–P bonds can occasionally occur at stitching junctions. To remedy this, we implemented an optional post-processing energy minimization step after coordinate writing. We used the OpenMM package [58] with the parmbsc1 force field [51] to run a short minimization in vacuum with a non-bonded interaction cut-off of 1.1 nm in a non-periodic system. Minimization was performed with a force tolerance of 500 kJ mol^−1^ nm^−1^. To quantify the effect of this procedure, we minimized 1000 HCG structures of the 33 bp sequence used in Ref. 34. We compared structures before and after minimization on distributions of O3^′^-P bonds across the structures as well as the RMSD of all heavy atoms in the chain. The minimization reliably corrected the long bonds while introducing minimal overall changes to the structure (Supporting Fig. 2). All analysis presented in the main text was performed before energy minimization. Thus, the minimization is only required if MD simulations are performed downstream, where energy minimizations are part of standard procedure.

### Efficient implementation of HCG-DNA enables atomistic representations on genome scale

Our new implementation of HCG employs a parallel Monte Carlo (MC) algorithm with a memory-efficient representation of atomic positions. Unlike previous implementations, we do not explicitly generate the complete set of atomic coordinates at every HCG level. Instead, we store recursively composed rigid-body transformations (**R**, **t**) that map each sampled fragment into its position in the full-length DNA polymer molecule (Supporting Fig. 3), where **R** ∈ SO(3) is a 3 × 3 rotation matrix and **t** ∈ R^3^ is a 3D vector, i.e., the pair is an element of the special Euclidean group of rigid-body transformations, (**R**, **t**) ∈ SE(3) anchored to the first fragment in the chain (Supporting Fig. 3). This approach is memory-efficient, since each fragment stitching attempt requires only the transformation operation and the 3D atomic positions of two MD fragments to be kept in memory. Atomic coordinates for the complete molecule are generated only when the final structures are written.

With this implementation, we generated atomically resolved DNA ensembles for sequences up to 1 Mbp (10^6^ bp) at close-to-ideal *L* log^2^ *L* scaling in computational cost (Supporting Fig. 4). To characterize computational scaling, we measured wall-clock times for the assembly of ensembles of 500 structures for sequences ranging from 2 kbp to 1 Mbp. The new implementation yields 500 structures for 10-kbp sequences in 120 min wall-clock time on a single node with 2×28 Intel(R) Xeon(R) Platinum 8280 CPU cores. For the 1-Mbp system, 500 structures were generated within ≈110 hours on 4 compute nodes. This extends HCG-DNA ensemble generation to DNA lengths approaching the genome sizes of minimal cells [59]. The implementation is parallelized also across compute nodes, with good scaling up to ∼4 nodes (Supporting Fig. 4B). For ensemble generation, trivial parallelization is possible using different random seeds.

### Comparison of HCG-DNA with atomistic simulations and experimental measurements

All analyses below were performed on ensembles of 10,000 HCG-DNA structures and used the MD-Analysis python package [60] unless stated otherwise.

#### Comparison of HCG-DNA to atomistic MD simulations

We generated ensembles of the same 33 bp sequence used in an independent previously reported 5-*µ*s atomistic MD simulation [34] (5^′^-GAGATGCTAACCCTGATCGCTGATTCCTTGGAC-3^′^) and computed helical parameters (shift, slide, rise, tilt, roll, and twist), excluding 3 terminal base pairs at each end to exclude contributions from fraying events. We did the same for a randomly generated 33 bp sequence (5^′^-AGCGGCGAGCGGGAGCCATTCGTTTACGTTAAT-3^′^) to further compare the sequence dependent properties along the helix (Fig. 2 and Supporting Figs. 5 and 6).

**Figure 2:**
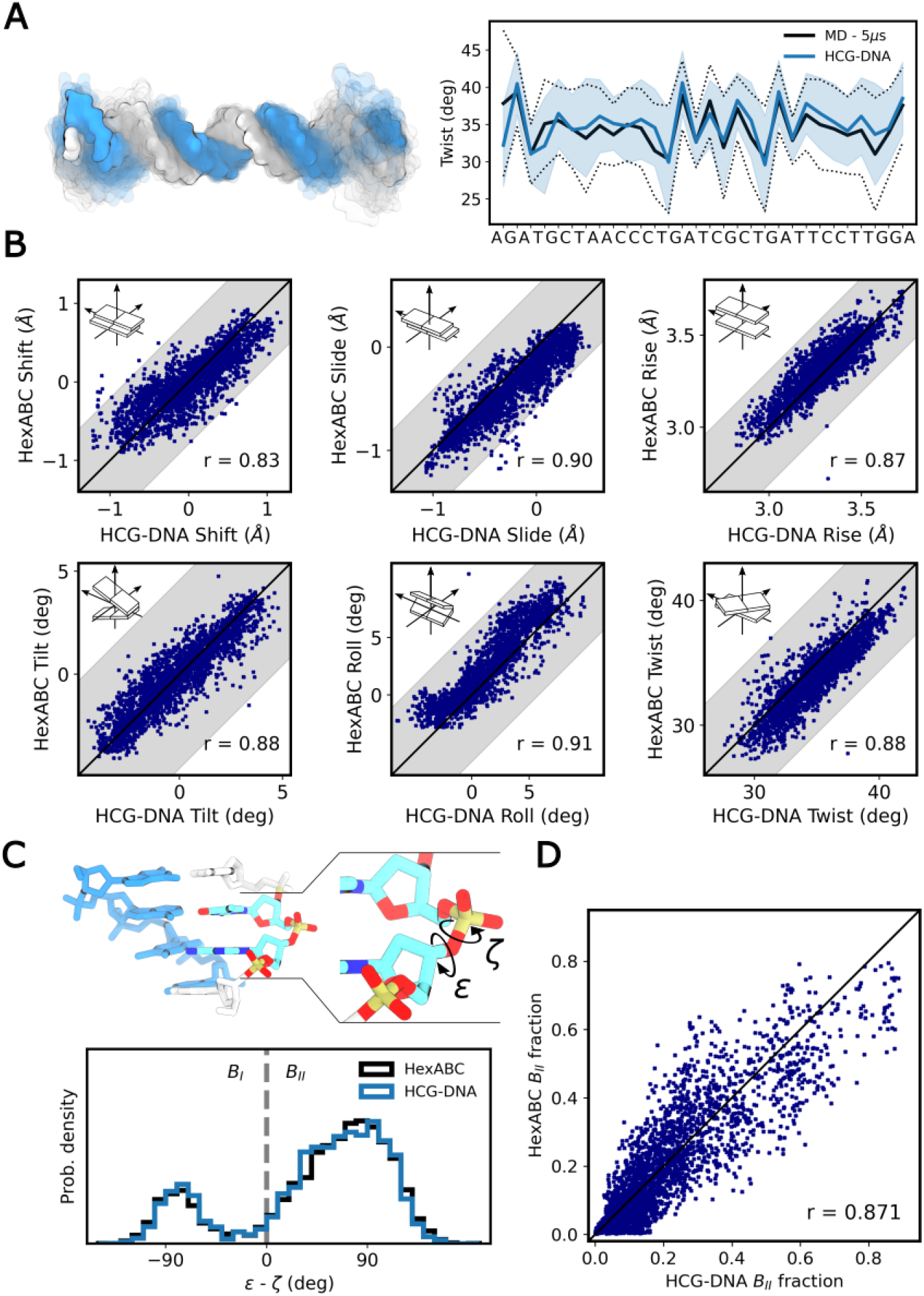
HCG-DNA preserves sequence-dependent DNA conformational statistics. (**A**) HCG-DNA ensemble for a 33 bp DNA duplex compared with a 5-*µ*s atomistic MD simulation. Left, representative HCG-DNA conformations. Right, position-resolved mean twist (solid lines) and standard deviation (shading/dashed lines) along the duplex for MD (black) and HCG-DNA (blue). (**B**) Sequence-dependent bp parameters for each base-pair step present in the hexABC MD simulation data set [33]. Each point compares the ensemble mean obtained from HCG-DNA with the mean from atomistic MD; Pearson correlation coefficients are shown. Grey regions indicate the average standard deviation of the corresponding MD distributions. (**C**,**D**) Sequence-dependent backbone conformations *B_I_*(*ε* − *ζ* ≤ 0^◦^) and *B_II_*(*ε* − *ζ >* 0^◦^) states defined from the backbone dihedrals *ε* and *ζ*. Bottom, *B_I_, B_II_* distributions for MD and HCG-DNA of the 5^′^-CTCpGCG-3^′^ hexanucleotide; (**D**) *B_II_* populations across the 2080 sequences of the hexABC data set (*r* = 0.871). *Alt text:* Graphs and molecular visualizations labeled A to D. A shows overlayed molecular visualizations on the left with one structure at full opacity and others at low opacity. The right side shows the twist per base-pair step with the sequence indicated on the x-axis. Lines and shaded areas line up closely for MD and HCG-DNA. B shows six correlation plots for shift, slide, rise, tilt, roll and twist listed left-to-right and top-to-bottom. Pictograms illustrate each parameter. Pearson correlation values are shown to lie between 0.83 and 0.91. C shows a molecular visualization of a tetranucleotide duplex with a zoom-in showing epsilon and zeta dihedral angles. Below that it shows histograms of epsilon minus zeta which line up closely between hexABC and HCG-DNA. D shows a correlation plot of B-2 fractions between HCG-DNA and hexABC.

For analysis across the full sequence space we built all 136 unique tetranucleotide sequences of ds-DNA, each embedded in a decamer (5^′^-GCGX_1_X_2_X_3_X_4_CGC-3^′^) to ensure that the central base-pair step (X_2_X_3_) enters into the stitched fragment pair (Fig. 1). We similarly embedded each tetranucleotide into an octamer (5^′^-CGX_1_X_2_X_3_X_4_CG-3^′^) to ensure the central base-pair step is inherited from the MD simulation. For each sequence context, we calculated the six helical parameters and the backbone dihedral angles *ε* and *ζ* at the central step and compared them with the corresponding values from the atomistic fragment simulations described above (Supporting Figs. 7 and 8).

For independent validation against longer sequence contexts, we performed a similar analysis using the hexABC dataset [33]. We generated ensembles of all unique 2080 DNA hexamers by creating the 190 20-bp DNA sequences in the hexABC set and calculated the helical parameters along the chain excluding the 3 terminal base pairs at each end. For each parameter, we compared ensemble-averaged values at corresponding base-pair steps between HCG-DNA and hexABC and quantified agreement using Pearson correlation coefficients (Fig. 2B). We additionally calculated *ε* and *ζ*, classified backbone conformations as *B_I_* (*ε* − *ζ* ≤ 0^◦^) or *B_II_* (*ε* − *ζ >* 0^◦^), and compared *B_II_* populations between the two ensembles (Fig. 2C,D). All analyses of helical descriptors and backbone dihedral angles used the Curves+ software [61].

#### Comparison of HCG-DNA to PELDOR experiments

We measured cross-strand distances in HCG-DNA ensembles of two 20 bp sequences (5^′^-GTCAGTCGCGCGCGCGCATC-3^′^, 5^′^-GTCAGTGCGCGCGCGCGATC-3^′^) all interspin distances reported in Fig. 1 of Ref. [30]. To approximate the position of the rigid cytosine spin label, we extrapolated the vector connecting the centers of the N1–C2 and C4–C5 bonds to a length of 0.77 nm, following the geometry used in Ref. [30]. For each labeled pair, distances were calculated for every conformation in the HCG-DNA ensemble, and the resulting mean and standard deviation were compared directly with the corresponding PELDOR measurements.

*Comparison of HCG-DNA to gene-size experiments:* We generated ensembles of the 4,361 bp linearized pBR322 plasmid (GenBank accession J01749.1 [62]) and compared them with the recent AFM measurements of Ref. [63]. End-to-end distances were calculated between the base-pair center of mass at either end of the helix. Following the analysis used for the experimental data, we fit the distribution obtained from the 10,000 HCG-DNA structures to the Gaussian chain probability distribution for the end-to-end distance *R_ee_* [64]:

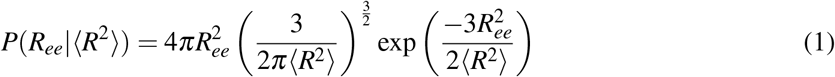

with ⟨*R*^2^⟩ as the fit parameter.

#### Persistence length from HCG-DNA ensembles

The persistence length *L_p_* quantifies the length scale at which correlations in monomer orientations persist along a polymer chain. We estimated *L_p_*for HCG-DNA ensembles of the 48,502 bp *λ*-phage genome (GenBank accession J02459 [62]) by calculating the dot product between base-pair normals *n̂_i_*and *n̂_j_*. For every structure, we scanned the chain with sliding windows of size *j* − *i* 1 to 500, then averaged the values for each window size. To obtain the persistence length, we again averaged across all 10,000 structures and fit Eq. 2, where ⟨*l*_0_⟩ = 0.34 nm is the average distance between base pairs, and *L_p_* is the persistence length:

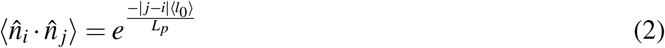

We estimated the uncertainty of *L_p_* as the standard deviation of fits of individual chains (Supporting Fig. 9).

#### Crookedness analysis

We measured the crookedness *β* in HCG ensembles of poly-A and poly-G 18-mers (5^′^-CGCGAAAAAAAAAACGCG-3^′^, 5^′^-CGCGGGGGGGGGGGCGCG-3^′^) used by Marin-Gonzalez et al. [65] defined as:

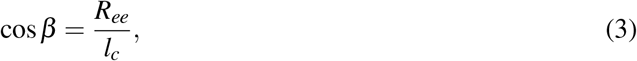

where *R_ee_* and *l_c_* represent the end-to-end distance and contour length, respectively. We calculated *R_ee_*and *l_c_* from base-pair center of masses, where *R_ee_*is the distance between the first and last base pairs and *l_c_* is the contour length of the centerline defined as the sum of all distances between subsequent base pairs (Supporting Fig. 10).

### Modeling linker DNA between nucleosomes with fixed positions and orientations measured by cryo-ET

We modeled nucleosome linkers in a cluster of 60 nucleosomes with 26 predicted linkers as part of the data published in Ref. 26. As a pool of conformers to screen for possible linker configurations, we generated a library spanning lengths from 16 to 106 bp including three additional base pairs for overlap with the nucleosomal DNA on each side. Since we do not have sequence-specificity in the alignment here, we chose 3 base pairs to match the overall helical structure. For each length, we generated ensembles of 1000 structures each for 100 random DNA sequences with 40% GC content on average.

To initialize the linker modeling, we positioned a nucleosome structure (PDB ID: pdb_00007dbp) at the positions and orientations obtained from subtomogram averaging. For each pair of connected nucleosomes, we screened the full library by aligning the three terminal base-pair backbones at each end to the corresponding nucleosomal DNA entry/exit points. We retained structures with an alignment RMSD below 2.5 Å and without steric clashes with the nucleosome or H1 histone. To assemble the full system, we randomly sampled from the shortest-length candidate sub-ensembles containing at least 20 structures until we obtained a clash-free combination. For nucleosomes without predicted connections, we only considered the histone proteins and the core 147 DNA base pairs during the clash check.

### Chromatin modeling with nucleosomes positioned on the sequence

#### Top-down

We generated conformational ensembles of chromatin by first generating linker ensembles of the target length and sequence. We then performed a second round of HCG in which linker-DNA and nucleosome conformations were treated as alternating fragments and assembled using the same alignment and clash-rejection procedure described above. We overlapped the ends of the linkers with base pairs 1 and 2 (entry site) or 146 and 147 (exit site) of the nucleosome. Minimum RMSD alignment and clash rejection were done as before. Clash rejection here included the histone protein heavy atoms for the close-range check with 2 Å cut-off and histone *C_α_* atoms for the long-range check with 4 Å cut-off. As nucleosome ensembles, we used the published MD simulation trajectory (15 *µ*s, 150 frames uniformly spaced) containing histone tails by Armeev et al. [66]. To facilitate alignment with the nucleosome entry and exit sites, we generated linker ensembles with two additional base pairs that matched with the two terminal base pairs of the nucleosome on each end for overlap. For tails, the adapter sequence was only applied to the overlapping end. Overlapping base pairs were discarded after assembly as done previously.

We generated ensembles of 100 structures of locus 58,110,000 to 58,210,000 on mouse chromosome 18 using nucleosome positioning data as provided in the public repository from Ref. 46. Assuming a length of 147 bp for nucleosome DNA, we excluded overlapping nucleosomes with a minimum linker length of 1 bp, yielding 305 nucleosomes. We generated linker ensembles to match the sequence of the same locus from the GRCm39 assembly.

#### Bottom-up

For analysis of 12-nucleosome arrays we generated ensembles of 1000 structures of randomly generated linker sequences with 40% GC content at effective lengths 1 to 50. The reported linker length excludes the base pairs added for overlap during assembly excluding the overlapping base pairs. For each length, we then generated 1000 structures of the chromatin chain with 12 nucleosomes, 11 linkers, and two tails. We measured the compaction of chromatin structures via the radius of gyration from positions of phosphorus atoms. For analysis of relative orientations of nucleosomes *i* and *i* + 2, we followed the analysis by Chen et al. [67]. We classified a pair as face-to-face, side-to-side, or face-to-side based on the acute angles (0-90^◦^) between nucleosome plane normals and the vector connecting the center of mass of the two nucleosomes (Fig. 4D). For face-to-face both normal vectors have to have an angle of *<* 45^◦^ to the connecting vector. For side-to-side both normal vectors need to be *>* 45^◦^ to the connecting vector. Face-to-side classification applies if one normal vector is *>* 45^◦^ and the other is *<* 45^◦^ degrees to the connecting vector. We then measured the fraction of all *i*, *i* + 2 nucleosome pairs that belong to each class and took the average and standard deviation across the ensemble. We additionally measured the average distance between *i*, *i* + 2 pairs and calculated the standard deviation across the ensemble.

To determine how incorporation of a DNA-bending transcription factor reshapes the conformational ensemble of a longer DNA molecule, we modeled the transcription factor LEF1-bound DNA using the 12 structures in the NMR ensemble of the LEF1 HMG domain (PDB ID pdb_00002lef [68]) in the HCG-DNA assembly of the corresponding DNA segment. We generated ensembles of 1000 structures of 200 bp duplexes containing 0, 1, 2 and 3 copies of the LEF1 HMG domain and the 15 bp included in the structure. The remaining sequence was randomly generated. Protein fragments were distributed as evenly as possible along the 200-bp duplex. For one bound protein, the 15-bp fragment was placed centrally, with flanking DNA segments of 93 bp and 92 bp at the 5^′^ and 3^′^ ends, respectively. For two bound proteins, we used a 57-bp internal linker and flanking segments of 57 bp (5^′^) and 56 bp (3^′^). For three bound proteins, we used two 39-bp internal linkers and flanking segments of 39 bp (5^′^) and 38 bp (3^′^). We measured the radius of gyration of the ensembles from positions of phosphorus atoms.

## Results

### Hierarchical assembly of double-stranded DNA

DNA structure and mechanics span orders of magnitude in length, from sequence-dependent molecular fluctuations to megabase-scale genome organization confined by the nucleus and the cell. Bridging these scales within a common molecular description remains challenging both experimentally and computationally. HCG-DNA addresses this challenge by constructing ensembles of DNA with atomistic resolution and defined sequence (Fig. 1). We first generated extensive atomistic MD libraries covering all 136 unique tetranucleotide sequence contexts and mapped a target sequence onto the corresponding overlapping fragment ensembles (see Methods and Supporting Fig. 1). In HCG-DNA, conformations sampled from these libraries are assembled hierarchically [48, 49]: overlapping coordinates are aligned, sterically incompatible combinations are rejected, and compatible subchains are recursively merged into progressively longer molecules. Repeating this procedure generates full-length structural ensembles while retaining atomic resolution throughout. HCG-DNA therefore propagates sequence-dependent molecular conformations from short DNA fragments to kilobase- and megabase-scale polymers and, by incorporating nucleosome-bound DNA, to atomistic chromatin ensembles.

HCG-DNA was designed to scale efficiently to large systems. Once the atomistic fragment libraries are generated, ensemble members can be sampled independently in parallel without rerunning explicit-solvent MD for each target sequence. A memory-efficient implementation further allows long DNA molecules to be assembled without storing complete polymer ensembles in memory, enabling rapid generation of atomically resolved structures from short duplexes to megabase-scale DNA (see details in Methods and Supporting Methods). This makes molecular-resolved sampling accessible at DNA lengths far beyond those tractable by direct atomistic simulation.

### HCG recovers sequence-dependent DNA ensembles from atomistic simulations

To assess whether hierarchical assembly preserves the sequence-dependent conformational information encoded in atomistic DNA simulations, we benchmarked HCG-DNA against a previously reported 5 *µ*s long, atomistic MD simulation of a 33 bp DNA duplex [34]. Across all six base-pair-step parameters (shift, slide, rise, tilt, roll, and twist), HCG-DNA reproduced the position-resolved means and standard deviations obtained from direct MD (Fig. 2A and Supporting Fig. 5). To test this systematically across all tetranucleotide sequence contexts, we embedded each of the 136 unique tetranucleotide contexts within octamer or decamer sequences (Methods), generated 10,000 HCG-DNA conformations for each sequence, and compared the six helical parameters of the central base-pair step with the corresponding atomistic fragment simulations. HCG-DNA recovered the sequence-dependent variation of all six parameters both when the central step was inherited directly from an MD fragment and when it was formed across a stitching junction (Supporting Fig. 7). Averaging over these two cases, as expected across assembled chains, yielded Pearson correlations of *r* = 0.94–0.98 across the six parameters.

We then challenged HCG-DNA against the recently reported hexABC dataset, which samples all 2,080 unique hexanucleotide contexts within 190 independently simulated 20 bp duplexes [33], albeit with different water and ion force-fields. We reconstructed the same 190 sequences using HCG-DNA and compared their sequence-dependent helical properties with the long atomistic simulations. Despite HCG-DNA being constructed solely from tetranucleotide libraries simulated with different simulation parameters and sequence contexts, HCG-DNA captured the sequence-dependent variation across all 2,080 hexanucleotide contexts, with Pearson correlations of *r* = 0.83–0.91 for the six base-pair-step parameters (Fig. 2B). Agreement with the hexABC reference extended beyond helical geometry to the major *B_I_*and *B_II_* conformational substates of the B-DNA backbone [69, 70]. Their populations, classified from the coupled *ε* and *ζ* backbone dihedrals, were similarly captured across the hexanucleotide contexts, with HCG-DNA and atomistic MD yielding correlated *B_II_* populations across sequence space (Pearson *r* = 0.871; Fig. 2C,D).

Furthermore, at the level of DNA shape, HCG-DNA also reproduced the previously reported difference in DNA crookedness between poly-A and poly-G duplexes (Supporting Fig. 10) [65]. Together, these comparisons show that hierarchical assembly preserves sequence-dependent molecular conformations and their effects on short DNA structures.

### Local molecular statistics predict DNA structure and mechanics across scales

Having established that hierarchical assembly preserves sequence-dependent molecular statistics, we asked whether the resulting ensembles reproduce independent experimental observables across scales. We first compared HCG-DNA with previously published PELDOR measurements that resolve intramolecular distance distributions with high precision for two 20 bp DNA duplexes [30]. For each sequence, we generated HCG-DNA ensembles and calculated intermolecular distances with the spin-label positions extrapolated from the cytosine nucleobase plane (Methods). Without any fitting, the HCG-DNA distributions closely reproduced the PELDOR measurements and were comparable to those obtained from direct atomistic MD using parmbsc1 [51] (Fig. 3A). The largest deviations occurred at the shortest distances, near the ≈ 18 Å lower detection limit of the experiment [30]. HCG-DNA therefore recovers an independent molecular-scale experimental observable directly from the conformational statistics encoded in its atomistic libraries.

**Figure 3:**
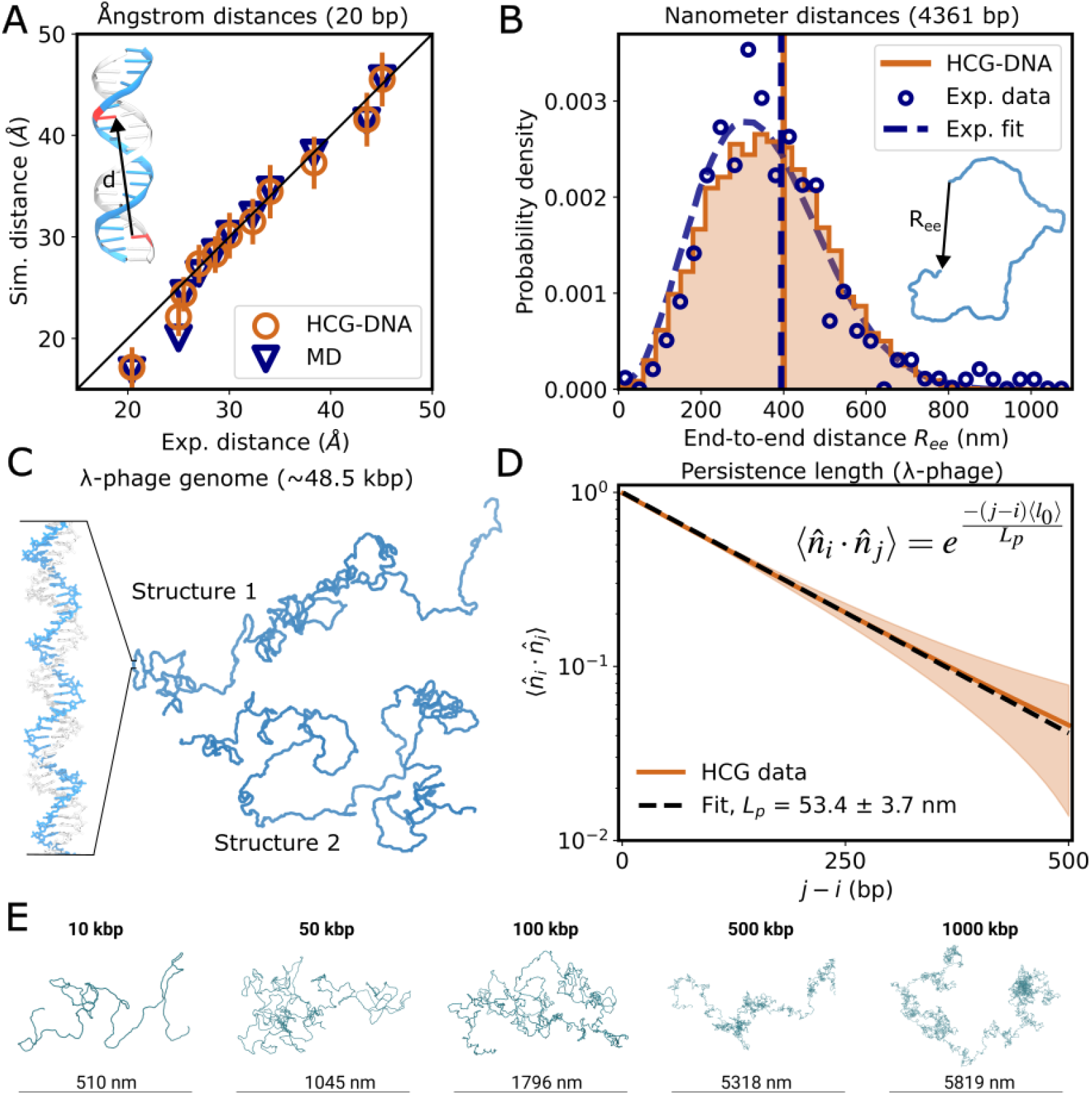
HCG-DNA connects molecular structure to genome-scale DNA mechanics. (**A**) Molecular-scale validation against PELDOR measurements for two 20-bp DNA duplexes. Experimentally measured cross-strand distances *d* are compared with distances obtained from HCG-DNA ensembles and published atomistic MD data [30]. Points show ensemble means and error bars indicate the corresponding standard deviations across the ensemble. (**B**) Kilobase-scale validation using the linearized 4,361-bp pBR322 plasmid. The HCG-DNA end-to-end distance distribution on top of the distribution measured by AFM on the same plasmid [63]. Lines show the a Gaussian-chain fit, and vertical lines indicate the root-mean-square end-to-end distances obtained from experiment and HCG-DNA. (**C**) Representative HCG-DNA conformations of the complete 48,502-bp *λ*-phage genome. (**D**) Orientational correlations between base-pair normals fitted with an exponential as a function of genomic separation ( *j* − *i*). Fitting this decay across 10,000 HCG-DNA conformations (10 representative examples are shown in Supporting Fig. 9) yields an average and standard deviation of the persistence length of *L_p_* = 53.4 ± 3.7 nm. Shading indicates the uncertainty associated with the 10,000 realizations. (**E**) Representative HCG-DNA conformations from 10-kbp to 1-Mbp illustrate the extension of the same atomistic representation across genomic length scales. *Alt text:* Graphs and molecular visualizations labeled A to E. A is plotting the correlation of distances from simulation on the y-axis and experiment on the x-axis. One such distance is shown on a molecular visualization of a DNA duplex. An exemplary distance Data from HCG-DNA and MD line up closely with each other and the experiment. B plots histograms of HCG-DNA and experimental data, which line up closely. A molecular visual- ization of a DNA model indicates how the end-to-end distance is measured. C shows two molecular visualizations with a zoom-in showing the atomic structure. D plots the decay of the correlation with increasing separation on the log-scale, giving a linear curve for the raw HCG-DNA data and the fit, which overlap exactly. E shows one molecular representation each for a 10, 50, 100, 500 and 1000 kbp duplex. Sizes range from 510 to 5819 nm.

We next tested whether the same molecular representation predicts collective DNA behavior at the kilobase scale. We generated an HCG-DNA ensemble of 10,000 structures of the linearized 4,361-bp pBR322 plasmid and compared its end-to-end distance distribution *R*_ee_ to recent AFM measurements [63]. The HCG-DNA end-to-end distribution closely overlapped with both the experimental histogram and the published three-dimensional Gaussian-chain fit used for the AFM data [63]. Applying the same Gaussian-chain description to the HCG-DNA ensemble (see Methods) yielded a root-mean-square end-to-end distance of 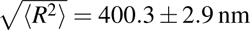 (fit ± s.e., Fig. 3B and Supporting Fig. 11), which is close to the moment estimate directly on the HCG-DNA data of 391.7 ± 1.4 nm (mean ± s.e.). This compares with 393 ± 9 nm measured by AFM (Fig. 3B) and 352 ± 24 nm independently predicted from magnetic-tweezers-derived polymer parameters [63] (mean ± s.e.). No polymer-scale dimension or elastic parameter was imposed during construction of the ensemble. Therefore, HCG-DNA predicts kilobase-scale polymer dimensions directly from local sequence-dependent conformational statistics propagated through hierarchical assembly.

We further asked whether molecular conformational statistics propagated by HCG-DNA are sufficient to recover the bending mechanics of genome-length DNA. We generated atomistic ensembles of the complete 48.5-kbp *λ*-phage genome (Fig. 3C), corresponding to the sequence unit used in a pioneering single-molecule force-extension experiments [71]. We determined the bending persistence length, *L_p_*, the characteristic distance over which DNAs tangent correlations decay (Methods). HCG-DNA yielded *L_p_* = 53.4 ± 3.7 nm (Fig. 3D), in excellent agreement with the range of values reported from multiple experiments [63, 72–74], including the worm-like-chain analysis of the same *λ*-DNA force–extension data [71]. Importantly, no persistence length or effective bending modulus was imposed during hierarchical assembly. Therefore, polymer bending mechanics at genomic-scale emerges from the molecular conformational statistics propagated from the atomistic fragment libraries.

Finally, we tested the computational efficiency of HCG-DNA assembly for increasingly large systems (Fig. 3E and Supporting Fig. 4). HCG-DNA scales approximately as *L* log^2^ *L* above lengths of about 6-kbp (Supporting Fig. 4). Our largest test case, the generation of an ensemble of 500 independent 1-Mbp conformations, each containing 4.1 × 10^7^ atoms (excluding hydrogen atoms), required 110 h on four dual-socket Intel Xeon Platinum 8280 nodes (224 cores; ≈ 50 core-hours per conformation).

### HCG-DNA connects experimental chromatin organization with atomistic ensembles

Chromatin organization is measured across scales, from genomic nucleosome maps to three-dimensional nucleosome arrangements resolved by cryo-ET [26] and fluorescence measurements in 3D [24]. Yet these observations provide limited insight into the underlying molecular structure. HCG-DNA connects these measurements to atomistic chromatin ensembles in two complementary directions. In a top-down approach, experimentally determined nucleosome positions, orientations or linker lengths constrain atomistic chromatin models (Fig. 4A-C). In a bottom-up approach, sequence-dependent linker DNA and nucleosome conformations are propagated into unconstrained higher-order ensembles, revealing how linker-DNA sequence and helical phasing shape nucleosome packing and chromatin compaction (Fig. 4D).

**Figure 4:**
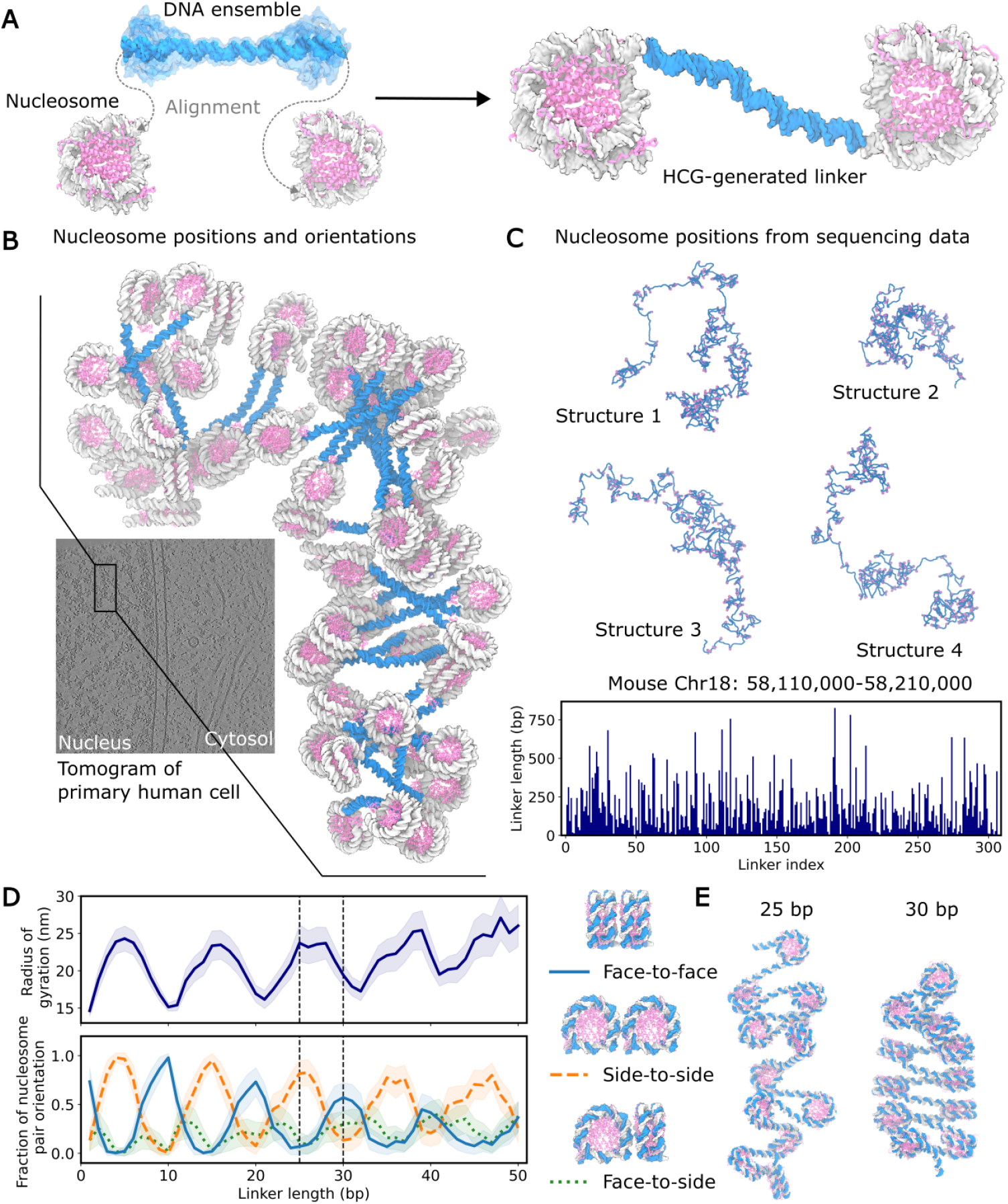
HCG-DNA connects experimental chromatin organization with atomistic ensembles and reveals linker-dependent packing. (**A**) HCG-DNA connects successive nucleosomes by sampling sequence-specific linker-DNA conformations that satisfy the entry and exit geometries of neighboring nucleosomes (see Methods). (**B**) Atomistic reconstruction of an in situ chromatin region containing 60 nucleosomes with 26 linkers using the approach from **A** and nucleosome positions and orientations determined by cryo-ET in a primary human cell. Data from Ref. 26. (**C**) Atomistic chromatin ensembles constrained by genomic nucleosome positions. Four representative conformations are shown for the locus 58,110,000 to 58,210,000 of the mouse chromosome 18 containing 306 linkers, with nucleosome positions derived from MNase-seq data from mouse embryonic stem cells [46]. Linker lengths across the locus are shown below. (**D**) Structural analysis of 12-nucleosome arrays as function of linker lengths from 1 to 50 bp. Top: Radius of gyration (mean ± s.d. shown as solid line and shading, respectively). Bottom: fraction of different orientations between nucleosomes *i* and *i* + 2. The corresponding distances are shown in Supporting Fig. 12. The legend on the right shows idealized cases of each of the orientation classes. (**E**) Representative structures with 25-bp and 30-bp linker lengths. *Alt text:* Graphs and molecular visualizations labeled A to E. a shows two nucleosomes and a set of overlayed DNA structures with two arrows labeled alignment pointing from the DNA ends to the nucleosome linker arms. An arrow points to the right, where a single DNA structure is placed to connect the nucleosomes. A shows an image of a tomographic slice with the nucleus on the left and cytosol on the right. A zoom into the nucleus shows a molecular representation of nucleosomes and linkers. C shows 4 molecular representations and a bar chart of linker lengths per linker. D shows curves of radius of gyration and fractions of nucleosome orientations over linker lengths showing a 10 base-pair periodicity. Local minima in radius of gyration line up with minima of side-to-side orientations and maxima of face-to-face orientations. E shows visualizations of two structures.

Cryo-ET can now provide the three-dimensional positions and orientations of individual nucleosomes inside cells together with predicted linker-DNA connections [26], thanks to optimized data collection and analysis protocols [75, 76]. Building on the data from Ref. 26, we used the experimentally determined nucleosome geometries as spatial constraints and HCG-DNA to sample linker conformations compatible with successive nucleosomes. This reconstructs fully atomistic chromatin models that satisfy the observed in situ architecture and can be used directly for molecular simulations (Fig. 4A,B).

A complementary top-down route starts from genomic nucleosome maps, which define nucleosome positions along the DNA sequence while leaving their three-dimensional organization unconstrained. Using MNase data for mouse chromosome 18 locus 58,110,000 to 58,210,000 [46], we placed nucleosomes at the experimentally inferred positions along the sequence and sampled the corresponding sequence-specific linker ensembles with HCG-DNA. For the dynamic nucleosome conformations, we used a publicly available long atomistic MD trajectory [66]. The resulting ensembles combine experimentally defined nucleosome placement with molecular heterogeneity in both linker DNA and nucleosome structure (Fig. 4C).

We next reversed the direction of inference and asked which features of chromatin organization emerge from molecular DNA and nucleosome conformations alone. For this, we generated unconstrained HCG-chromatin ensembles with defined linker lengths. Varying linker length produced a pronounced periodic modulation of chromatin dimensions and nucleosome packing, consistent with the helical periodicity of DNA (Fig. 4D). Linkers differing by 5 bp, approximately half a DNA helical turn, favored distinct relative nucleosome orientations and packing arrangements, as illustrated by the markedly different structures formed with 25- and 30 bp linkers. These trends recapitulate recent experimental and simulation studies showing that linker-DNA phasing controls nucleosome stacking and higher-order chromatin organization [43, 67, 77, 78]. HCG-DNA therefore propagates local DNA geometry into mesoscale chromatin architecture, revealing how helical phasing encoded at the linker level can tune nucleosome packing and global chromatin compaction.

Beyond nucleosome organization, HCG-DNA can propagate local structural perturbations introduced by DNA-binding proteins into larger-scale changes in DNA conformation. To test this, we incorporated increasing numbers of the bend-inducing transcription factor LEF-1 into 200-bp HCG-DNA chains (Supporting Fig. 13). LEF-1 introduces a pronounced local DNA bend and has been implicated in chromatin reorganization and gene looping [68, 79]. Increasing LEF-1 occupancy progressively shifted the radius-of-gyration distributions toward smaller values and broadened the accessible conformational ensemble, producing markedly more compact structures than unbound DNA. Thus, local protein-induced DNA deformations can propagate into global changes in DNA organization.

Together, these top-down and bottom-up analyses show that HCG-DNA can both reconstruct experimentally observed chromatin organization at atomic resolution and reveal how local DNA geometry propagates into higher-order chromatin structure.

## Discussion

Generating atomistically resolved ensembles of DNA and chromatin across physiologically relevant length scales remains challenging. Here, we found that local atomistic DNA conformations retain quantitative information when propagated across several orders of magnitude in length. HCG-DNA preserves sequence-dependent helical and backbone properties (Fig. 2 and Supporting Figs. 5-8) observed in long atomistic simulations [34] and in the independent hexABC [33] (Fig. 2). Remaining deviations may reflect both differences in the water and ion force fields used here and for hexABC and sequence-context effects beyond the tetranucleotide level. The HCG-DNA ensembles further reproduce experimental molecular distance distributions [30] and polymer-scale properties [63, 71–74]. HCG-DNA predicts a persistence length of *L_p_* = 53.4 nm, without fitting a bending modulus or other polymer-scale elastic parameter in excellent agreement with experimental reports [71–74]. Similarly, the dimensions of kilobase DNA emerge from the same molecular representation (Fig. 3). Thus, molecular conformational statistics sampled at the tetranucleotide level is sufficient to recover properties that only become apparent at much larger length scales.

HCG-DNA produces atomistic conformational ensembles that extend across scales. Long-chain ensembles are assembled from sampled molecular conformations, while the hierarchical construction preserves the statistical weights of compatible structures under the geometric and steric constraints of the model [47, 48]. In contrast to approaches that transfer atomistic information through fitted reduced representations [38, 40, 42], no intermediate-resolution Hamiltonian is fitted and no back-mapping step is required. Importantly, HCG-DNA is not parameterized against polymer-scale observables. Instead, polymer dimensions, persistence length, and chromatin compaction emerge from local structure and fluctuations.

HCG-DNA furthermore links spatial information from experiments to molecular structural ensembles. We have shown before how HCG-DNA together with nucleosome positions and orientations determined by cryo-ET generate atomic structures of in situ human chromatin patch (see Ref. 26 and Fig. 4A,B). Additionally, using genomic nucleosome maps to define nucleosome placement along the sequence, HCG builds ensembles of a gene locus Fig. 4C). Conversely, bottom-up chromatin ensembles show that single-base-pair changes in linker length produce changes in nucleosome orientation that affect packing and chromatin compaction, revealing how local DNA geometry translates into higher-order organization without imposing a predefined chromatin fiber (Fig. 4D,E). Recent OpenCGChromatin simulations likewise reproduce the oscillatory dependence of chromatin packing on linker length using an explicit coarse-grained energetic model that includes DNA, nucleosome, and histone-tail interactions [43]. The emergence of the same periodicity in HCG-DNA without fitting chromatin-scale compaction highlights the contribution of linker-DNA geometry to higher-order organization.

The HCG assembly procedure readily incorporates the increasingly rich spatial information from cryo-ET, FISH, chromatin tracing or other imaging approaches to constrain progressively larger genomic regions [24–26, 80, 81]. Using robotics approaches [82], HCG can be adapted to target endpoints and trace tracks [83]. HCG-generated ensembles can also serve as molecular representations against which Förster resonance energy transfer (FRET), small-angle X-ray and neutron scattering (SAXS/SANS) or other ensemble-sensitive experiments are interpreted. Furthermore, HCG-generated structures provide starting configurations for coarse-grained or atomistic simulations [42, 43, 46, 84] when dynamics is required.

As possible issues, HCG-DNA inherits the sequence context and force-field dependence of its fragment libraries. Moreover, at larger scales, long-range electrostatics, DNA topology and supercoiling, nucleosome–nucleosome interactions, histone tails and molecular crowding will require additional physical descriptions. HCG-DNA provides atomistic ensembles onto which such interactions or experimental spatial constraints can be added. More broadly, our results establish that molecular conformational ensembles can be propagated directly from local DNA structure to polymer and chromatin scales, providing a route toward experimentally constrained molecular models of genome organization for biology and machine learning.

## Data availability

The code and the tetranucleotide fragment libraries will be publicly released upon publication of this article.

The DNA sequences analyzed in this work are publicly available from GenBank under accession numbers J01749.1 (pBR322) and J02459 (*λ*-phage), and the mouse genomic sequence was obtained from the GRCm39 reference assembly. Previously published simulation and experimental datasets used for validation are available from the original publications cited in the text.

## Author contributions statement

J.B.: Conceptualization, Data curation, Formal analysis, Investigation, Methodology, Software, Validation, Visualization, Writing - original draft, Writing - review & editing. S.C.-L.: Conceptualization, Data curation, Formal analysis, Investigation, Methodology, Project administration, Supervision, Validation, Writing - original draft, Writing - review & editing. T.H.: Conceptualization, Data curation, Formal analysis, Investigation, Methodology, Software, Validation, Writing - original draft, review & editing. J.D. Conceptualization, Writing - review & editing. G.H. Conceptualization, Funding acquisition, Investigation, Project administration, Resources, Supervision, Writing - original draft, Writing - review & editing.

## Acknowledgments

We thank Jürgen Köfinger and Lisa M. Pietrek for helpful guidance and discussion. We thank the Max Planck Society for its support and the Max Planck Computing and Data Facility for computational resources. The authors used Claude Opus 5.5 to assist with data analysis scripts, code development and manuscript formatting. All output was reviewed and verified by the authors, who take full responsibility for the content.

## References

[1] Tom Misteli. The self-organizing genome: principles of genome architecture and function. Cell, 183:28–45, 2020. doi: 10.1016/j.cell.2020.09.014.

[2] Robin Weinmann, Lukas Frank, and Karsten Rippe. Approaches to characterize chromatin subcompartment organization in the cell nucleus. Curr. Opin. Struct. Biol., 83:102695, 2023. doi: 10.1016/j.sbi.2023.102695.

[3] Satoshi Fujii, Hidetoshi Kono, Shigeori Takenaka, Nobuhiro Go, and Akinori Sarai. Sequence-dependent DNA deformability studied using molecular dynamics simulations. Nucleic Acids Res., 35:6063–6074, 2007. doi: 10.1093/nar/gkm627.

[4] Remo Rohs, Sean M. West, Alona Sosinsky, Peng Liu, Richard S. Mann, and Barry Honig. The role of DNA shape in protein–DNA recognition. Nature, 461:1248–1253, 2009. doi: 10.1038/nature08473.

[5] Filip Lankaš, Jiřı Šponer, Pavel Hobza, and Jörg Langowski. Sequence-dependent elastic properties of DNA. J. Mol. Biol., 299:695–709, 2000.

[6] Pablo D Dans, Alberto Perez, Ignacio Faustino, Richard Lavery, and Modesto Orozco. Exploring polymorphisms in B-DNA helical conformations. Nucleic Acids Res., 40:10668–10678, 2012.

[7] Pablo Daniel Dans, Ignacio Faustino, Federica Battistini, Krystyna Zakrzewska, Richard Lavery, and Modesto Orozco. Unraveling the sequence-dependent polymorphic behavior of d(CpG) steps in B-DNA. Nucleic Acids Res., 42:11304–11320, 2014.

[8] Marco Pasi, John H Maddocks, David Beveridge, Thomas C Bishop, David A Case, Thomas Cheatham, III, Pablo D Dans, B Jayaram, Filip Lankas, Charles Laughton, et al. *µ*ABC: a systematic microsecond molecular dynamics study of tetranucleotide sequence effects in B-DNA. Nucleic Acids Res., 42:12272–12283, 2014.

[9] Sergio Cruz-León and Nadine Schwierz. Hofmeister series for metal-cation–RNA interactions: the interplay of binding affinity and exchange kinetics. Langmuir, 36:5979–5989, 2020. doi: 10.1021/acs.langmuir.0c00851.

[10] Sergio Cruz-León and Nadine Schwierz. RNA captures more cations than DNA: insights from molecular dynamics simulations. J. Phys. Chem. B, 126:8646–8654, 2022. doi: 10.1021/acs.jpcb.2c04488.

[11] PT Lowary and Jonathan Widom. New DNA sequence rules for high affinity binding to histone octamer and sequence-directed nucleosome positioning. J. Mol. Biol., 276:19–42, 1998.

[12] Eran Segal, Yvonne Fondufe-Mittendorf, Lingyi Chen, AnnChristine Thåström, Yair Field, Irene K Moore, Ji-Ping Z Wang, and Jonathan Widom. A genomic code for nucleosome positioning. Nature, 442:772–778, 2006.

[13] Noam Kaplan, Irene K Moore, Yvonne Fondufe-Mittendorf, Andrea J Gossett, Desiree Tillo, Yair Field, Emily M LeProust, Timothy R Hughes, Jason D Lieb, Jonathan Widom, et al. The DNA-encoded nucleosome organization of a eukaryotic genome. Nature, 458:362–366, 2009.

[14] Kevin Struhl and Eran Segal. Determinants of nucleosome positioning. Nat. Struct. Mol. Biol., 20:267–273, 2013. doi: 10.1038/nsmb.2506.

[15] Shuming Liu, Advait Athreya, Zhuohan Lao, and Bin Zhang. From nucleosomes to compartments: physicochemical interactions underlying chromatin organization. Annu. Rev. Biophys., 53, 2024.

[16] Joshua Moller and Juan J de Pablo. Bottom-up meets top-down: the crossroads of multiscale chromatin modeling. Biophys. J., 118:2057–2065, 2020.

[17] Jan Huertas, M Julia Maristany, Jose Ignacio Perez-Lopez, and Rosana Collepardo-Guevara. Pushing the experimental resolution boundary in chromatin fibre organisation using multiscale simulations. Curr. Opin. Genet. Dev., 98:102464, 2026.

[18] Guo-Cheng Yuan, Yuen-Jong Liu, Michael F. Dion, Michael D. Slack, Lani F. Wu, Steven J. Altschuler, and Oliver J. Rando. Genome-scale identification of nucleosome positions in *S. cerevisiae*. Science, 309: 626–630, 2005. doi: 10.1126/science.1112178.

[19] Erez Lieberman-Aiden, Nynke L. van Berkum, Louise Williams, Maxim Imakaev, Tobias Ragoczy, Agnes Telling, Ido Amit, Bryan R. Lajoie, Peter J. Sabo, Michael O. Dorschner, Richard Sandstrom, Bradley Bernstein, M. A. Bender, Mark Groudine, Andreas Gnirke, John Stamatoyannopoulos, Leonid A. Mirny, Eric S. Lander, and Job Dekker. Comprehensive mapping of long-range interactions reveals folding principles of the human genome. Science, 326:289–293, 2009. doi: 10.1126/science.1181369.

[20] Jason D. Buenrostro, Beijing Wu, Howard Y. Chang, and William J. Greenleaf. ATAC-seq: a method for assaying chromatin accessibility genome-wide. Curr. Protoc. Mol. Biol., 109:21.29.1–21.29.9, 2015. doi: 10.1002/0471142727.mb2129s109.

[21] Dustin E Schones, Kairong Cui, Suresh Cuddapah, Tae-Young Roh, Artem Barski, Zhibin Wang, Gang Wei, and Keji Zhao. Dynamic regulation of nucleosome positioning in the human genome. Cell, 132:887–898, 2008.

[22] Siyuan Wang, Jun-Han Su, Brian J Beliveau, Bogdan Bintu, Jeffrey R Moffitt, Chao-ting Wu, and Xiaowei Zhuang. Spatial organization of chromatin domains and compartments in single chromosomes. Science, 353: 598–602, 2016.

[23] Leslie J Mateo, Sedona E Murphy, Antonina Hafner, Isaac S Cinquini, Carly A Walker, and Alistair N Boettiger. Visualizing DNA folding and RNA in embryos at single-cell resolution. Nature, 568:49–54, 2019.

[24] K. S. Beckwith, Ø Ødegård-Fougner, N. R. Morero, C. Barton, F. Schueder, W. Tang, S. Alexander, J.-M. Peters, R. Jungmann, E. Birney, and J. Ellenberg. Nanoscale 3D DNA tracing in non-denatured cells resolves the cohesin-dependent loop architecture of the genome *in situ*. Nat. Commun., 16:6673, 2025. doi: 10.1038/s41467-025-61689-y.

[25] Zhen Hou, Frank Nightingale, Yanan Zhu, Craig MacGregor-Chatwin, and Peijun Zhang. Structure of native chromatin fibres revealed by cryo-ET *in situ*. Nat. Commun., 14:6324, 2023. doi: 10.1038/s41467-023-42072-1.

[26] Jan Philipp Kreysing, Sergio Cruz-León, Johannes Betz, Carlotta Penzo, Tomáš Majtner, Markus Schreiber, Beata Turoňová, Marina Lusic, Gerhard Hummer, and Martin Beck. Molecular architecture of heterochromatin at the nuclear periphery of primary human cells. Nat. Commun., 17:5844, 2026. doi: 10.1038/s41467-026-75087-5.

[27] Richard Wing, Horace Drew, Tsunehiro Takano, Chris Broka, Shoji Tanaka, Keiichi Itakura, and Richard E Dickerson. Crystal structure analysis of a complete turn of B-DNA. Nature, 287:755–758, 1980.

[28] Wilma K. Olson, Andrey A. Gorin, Xiang-Jun Lu, Lynette M. Hock, and Victor B. Zhurkin. DNA sequence-dependent deformability deduced from protein–DNA crystal complexes. Proc. Natl. Acad. Sci. U.S.A., 95: 11163–11168, 1998. doi: 10.1073/pnas.95.19.11163.

[29] John Kuszewski, Charles Schwieters, and G. Marius Clore. Improving the accuracy of NMR structures of DNA by means of a database potential of mean force describing base–base positional interactions. J. Am. Chem. Soc., 123:3903–3918, 2001. doi: 10.1021/ja010033u.

[30] Lukas S. Stelzl, Nicole Erlenbach, Marcel Heinz, Thomas F. Prisner, and Gerhard Hummer. Resolving the conformational dynamics of DNA with ångstrom resolution by pulsed electron–electron double resonance and molecular dynamics. J. Am. Chem. Soc., 139:11674–11677, 2017. doi: 10.1021/jacs.7b05363.

[31] H.G. Hansma, D.E. Laney, M. Bezanilla, R.L. Sinsheimer, and P.K. Hansma. Applications for atomic force microscopy of DNA. Biophys. J., 68:1672–1677, 1995. doi: 10.1016/S0006-3495(95)80343-7.

[32] Steven B. Smith, Laura Finzi, and Carlos Bustamante. Direct mechanical measurements of the elasticity of single DNA molecules by using magnetic beads. Science, 258:1122–1126, 1992. doi: 10.1126/science.1439819.

[33] Federica Battistini, Miłosz Wieczór, Adam Hospital, Marco Pasi, Juan Pablo Arcón, Israel Serrano-Chacón, Alba Sala, Subhamoy Deb, Agustín García-Doñate, Matthew Burman, Elliot W. Chan, Liwei Chang, Gabriela da Rosa, Jorge R. Espinosa, Gia Linh Hoang, Kazi A. Hossain, Michał Jurkowski, Romain Poupon, Rahul Sharma, Ran Sun, Thomas C. Bishop, Paolo Carloni, Thomas E. Cheatham, Rosana Collepardo-Guevara, Jacek Czub, Pablo D. Dans, Sarah A. Harris, Charles Laughton, Rodrigo Galindo-Murillo, John H. Mad-docks, Agnes Noy, Alberto Pérez, Daiva Petkevičiūtė-Gerlach, and Modesto Orozco. hexABC seeking the physical code of DNA. Nat. Commun., 17:8169, 2026. doi: 10.1038/s41467-026-74390-5.

[34] Sergio Cruz-León, Willem Vanderlinden, Peter Müller, Tobias Forster, Georgina Staudt, Yi-Yun Lin, Jan Lipfert, and Nadine Schwierz. Twisting DNA by salt. Nucleic Acids Res., 50:5726–5738, 2022. doi: 10.1093/nar/gkac445.

[35] Koen R Storm, Christian Wiebeler, Sergio Cruz-León, Caroline Körösy, Nadine Schwierz, and Jan Lipfert. DNA twist at high alkali ion concentrations: evidence against C-form DNA in solution. Nucleic Acids Res., 54:gkag192, 2026. doi: 10.1093/nar/gkag192.

[36] Petr Šulc, Flavio Romano, Thomas E. Ouldridge, Lorenzo Rovigatti, Jonathan P. K. Doye, and Ard A. Louis. Sequence-dependent thermodynamics of a coarse-grained DNA model. J. Chem. Phys., 137:135101, 2012. doi: 10.1063/1.4754132.

[37] Jaakko J. Uusitalo, Helgi I. Ingólfsson, Parisa Akhshi, D. Peter Tieleman, and Siewert J. Marrink. Martini coarse-grained force field: extension to DNA. J. Chem. Theory Comput., 11:3932–3945, 2015. doi: 10.1021/acs.jctc.5b00286.

[38] Jürgen Walther, Pablo D Dans, Alexandra Balaceanu, Adam Hospital, Genís Bayarri, and Modesto Orozco. A multi-modal coarse grained model of DNA flexibility mappable to the atomistic level. Nucleic Acids Res., 48:e29, 2020. doi: 10.1093/nar/gkaa015.

[39] Stephanie Portillo-Ledesma and Tamar Schlick. Bridging chromatin structure and function over a range of experimental spatial and temporal scales by molecular modeling. Wiley Interdiscip. Rev. Comput. Mol. Sci., 10:e1434, 2020. doi: 10.1002/wcms.1434.

[40] Rahul Sharma, Alessandro S. Patelli, Lennart De Bruin, and John H. Maddocks. cgNA+web: a visual interface to the cgNA+ sequence-dependent statistical mechanics model of double-stranded nucleic acids. J. Mol. Biol., 435:167978, 2023. doi: 10.1016/j.jmb.2023.167978.

[41] Sergio Cruz-León, Kara K. Grotz, and Nadine Schwierz. Extended magnesium and calcium force field parameters for accurate ion–nucleic acid interactions in biomolecular simulations. J. Chem. Phys., 154: 171102, 2021. doi: 10.1063/5.0048113.

[42] David Farré-Gil, Juan Pablo Arcon, Charles A Laughton, and Modesto Orozco. CGeNArate: a sequence-dependent coarse-grained model of DNA for accurate atomistic MD simulations of kb-long duplexes. Nucleic Acids Res., 52:6791–6801, 2024. doi: 10.1093/nar/gkae444.

[43] Kieran Russell, Yifang Chen, Jorge R. Espinosa, David Farré-Gil, Huabin Zhou, M. Julia Maristany, Jose Ignacio Perez-Lopez, Jan Huertas, Modesto Orozco, Michael K. Rosen, and Rosana Collepardo-Guevara. Near- atomistic simulations reveal the molecular principles that control chromatin structure and phase separation. Nat. Commun., 17:10532, 2026. doi: 10.1038/s41467-026-78050-6.

[44] Ivan Riveros and Bin Zhang. NEAT-DNA: a chemically accurate, sequence-dependent coarse-grained model for large-scale DNA simulations. J. Chem. Theory Comput., 22:3709–3719, 2026. doi: 10.1021/acs.jctc.5c01966.

[45] Rosana Collepardo-Guevara, Guillem Portella, Michele Vendruscolo, Daan Frenkel, Tamar Schlick, and Modesto Orozco. Chromatin unfolding by epigenetic modifications explained by dramatic impairment of internucleosome interactions: a multiscale computational study. J. Am. Chem. Soc., 137:10205–10215, 2015. doi: 10.1021/jacs.5b04086.

[46] Zilong Li and Tamar Schlick. Hi-BDiSCO: folding 3D mesoscale genome structures from Hi-C data using Brownian dynamics. Nucleic Acids Res., 52:583–599, 2024. doi: 10.1093/nar/gkad1121.

[47] Lisa M. Pietrek, Lukas S. Stelzl, and Gerhard Hummer. Structural ensembles of disordered proteins from hierarchical chain growth and simulation. Curr. Opin. Struct. Biol., 78:102501, 2023. doi: 10.1016/j.sbi.2022.102501.

[48] Lisa M. Pietrek, Lukas S. Stelzl, and Gerhard Hummer. Hierarchical ensembles of intrinsically disordered proteins at atomic resolution in molecular dynamics simulations. J. Chem. Theory Comput., 16:725–737, 2020.

[49] Lisa M. Pietrek, Lukas S. Stelzl, and Gerhard Hummer. Hierarchical assembly of single-stranded RNA. J. Chem. Theory Comput., 20:2246–2260, 2024. doi: 10.1021/acs.jctc.3c01049.

[50] Mark James Abraham, Teemu Murtola, Roland Schulz, Szilárd Páll, Jeremy C. Smith, Berk Hess, and Erik Lindahl. GROMACS: high performance molecular simulations through multi-level parallelism from laptops to supercomputers. SoftwareX, 1-2:19–25, 2015.

[51] Ivan Ivani, Pablo D. Dans, Agnes Noy, Alberto Pérez, Ignacio Faustino, Adam Hospital, Jürgen Walther, Pau Andrio, Ramon Goñi, Alexandra Balaceanu, Guillem Portella, Federica Battistini, Josep Lluis Gelpí, Carlos González, Michele Vendruscolo, Charles A. Laughton, Sarah A. Harris, David A. Case, and Modesto Orozco. Parmbsc1: a refined force field for DNA simulations. Nat. Methods, 13:55–58, 2016.

[52] Shavkat Mamatkulov and Nadine Schwierz. Force fields for monovalent and divalent metal cations in TIP3P water based on thermodynamic and kinetic properties. J. Chem. Phys., 148:074504, 2018.

[53] Kara K. Grotz and Nadine Schwierz. Optimized magnesium force field parameters for biomolecular simulations with accurate solvation, ion-binding, and water-exchange properties in SPC/E, TIP3P-fb, TIP4P/2005, TIP4P-Ew, and TIP4P-D. J. Chem. Theory Comput., 18:526–537, 2022.

[54] José Guadalupe Rosas Jiménez, Balázs Fábián, and Gerhard Hummer. Faster sampling in molecular dynamics simulations with TIP3P-F water. J. Chem. Theory Comput., 20:11068–11081, 2024.

[55] H. J. C. Berendsen, J. P. M. Postma, W. F. van Gunsteren, A. DiNola, and J. R. Haak. Molecular dynamics with coupling to an external bath. J. Chem. Phys., 81:3684–3690, 1984. doi: 10.1063/1.448118.

[56] Giovanni Bussi, Davide Donadio, and Michele Parrinello. Canonical sampling through velocity rescaling. J. Chem. Phys., 126:014101, 2007.

[57] Mattia Bernetti and Giovanni Bussi. Pressure control using stochastic cell rescaling. J. Chem. Phys., 153: 114107, 2020. doi: 10.1063/5.0020514.

[58] Peter Eastman, Raimondas Galvelis, Raúl P. Peláez, Charlles R. A. Abreu, Stephen E. Farr, Emilio Gallicchio, Anton Gorenko, Michael M. Henry, Frank Hu, Jing Huang, Andreas Krämer, Julien Michel, Joshua A. Mitchell, Vijay S. Pande, João PGLM Rodrigues, Jaime Rodriguez-Guerra, Andrew C. Simmonett, Sukrit Singh, Jason Swails, Philip Turner, Yuanqing Wang, Ivy Zhang, John D. Chodera, Gianni De Fabritiis, and Thomas E. Markland. OpenMM 8: molecular dynamics simulation with machine learning potentials. J. Phys. Chem. B, 128:109–116, 2024. doi: 10.1021/acs.jpcb.3c06662.

[59] Marian Breuer, Emmy E Earnest, Chuck Merryman, Kim S Wise, Lijie Sun, Michaela R Lynott, Clyde A Hutchison, Hamilton O Smith, John D Lapek, David J Gonzalez, Valérie de Crécy-Lagard, Drago Haas, Andrew D Hanson, Piyush Labhsetwar, John I Glass, and Zaida Luthey-Schulten. Essential metabolism for a minimal cell. eLife, 8:e36842, 2019.

[60] Naveen Michaud-Agrawal, Elizabeth J. Denning, Thomas B. Woolf, and Oliver Beckstein. MDAnalysis: a toolkit for the analysis of molecular dynamics simulations. J. Comput. Chem., 32:2319–2327, 2011.

[61] R. Lavery, M. Moakher, J. H. Maddocks, D. Petkeviciute, and K. Zakrzewska. Conformational analysis of nucleic acids revisited: Curves+. Nucleic Acids Res., 37:5917–5929, 2009.

[62] Dennis A. Benson, Mark Cavanaugh, Karen Clark, Ilene Karsch-Mizrachi, David J. Lipman, James Ostell, and Eric W. Sayers. GenBank. Nucleic Acids Res., 41:D36–D42, 2013. doi: 10.1093/nar/gks1195.

[63] Koen R. Storm, Caroline Körösy, Enrico Skoruppa, Stefanie D. Pritzl, Pauline J. Kolbeck, Willem Vanderlinden, Helmut Schiessel, and Jan Lipfert. The effects of DMSO on DNA conformations and mechanics. Biophys. J., 124:2639–2654, 2025.

[64] Michael Rubinstein and Ralph H Colby. Polymer Physics. Oxford University Press, Oxford, 2003. doi: 10.1093/oso/9780198520597.001.0001.

[65] Alberto Marin-Gonzalez, J. G. Vilhena, Fernando Moreno-Herrero, and Ruben Perez. DNA crookedness regulates DNA mechanical properties at short length scales. Phys. Rev. Lett., 122:048102, 2019. doi: 10.1103/PhysRevLett.122.048102.

[66] Grigoriy A. Armeev, Anastasiia S. Kniazeva, Galina A. Komarova, Mikhail P. Kirpichnikov, and Alexey K. Shaytan. Histone dynamics mediate DNA unwrapping and sliding in nucleosomes. Nat. Commun., 12:2387, 2021.

[67] Lifeng Chen, M. Julia Maristany, Stephen E. Farr, Jinyue Luo, Bryan A. Gibson, Lynda K. Doolittle, Jorge R. Espinosa, Jan Huertas, Sy Redding, Rosana Collepardo-Guevara, and Michael K. Rosen. Nucleosome spacing can fine-tune higher-order chromatin assembly. Nat. Commun., 16:6315, 2025.

[68] John J. Love, Xiang Li, David A. Case, Klaus Giese, Rudolf Grosschedl, and Peter E. Wright. Structural basis for DNA bending by the architectural transcription factor LEF-1. Nature, 376:791–795, 1995. doi: 10.1038/376791a0.

[69] M.N. Westwood, C.C. Johnson, Nathan A. Oyler, and Gary A. Meints. Kinetics and thermodynamics of B_I_– B_II_ interconversion altered by T:G mismatches in DNA. Biophys. J., 121:1691–1703, 2022. doi: 10.1016/j.bpj.2022.03.031.

[70] D. Djuranovic and B. Hartmann. DNA fine structure and dynamics in crystals and in solution: the impact of B_I_/B_II_ backbone conformations. Biopolymers, 73:356–368, 2004. doi: 10.1002/bip.10528.

[71] C. Bustamante, J. F. Marko, E. D. Siggia, and S. Smith. Entropic elasticity of *λ*-phage DNA. Science, 265: 1599–1600, 1994. doi: 10.1126/science.8079175.

[72] Jan Lipfert, Xiaomin Hao, and Nynke H. Dekker. Quantitative modeling and optimization of magnetic tweezers. Biophys. J., 96:5040–5049, 2009. doi: 10.1016/j.bpj.2009.03.055.

[73] C. Bouchiat, M.D. Wang, J.-F. Allemand, T. Strick, S.M. Block, and V. Croquette. Estimating the persistence length of a worm-like chain molecule from force-extension measurements. Biophys. J., 76:409–413, 1999. doi: 10.1016/S0006-3495(99)77207-3.

[74] Jan Lipfert, Sven Klijnhout, and Nynke H. Dekker. Torsional sensing of small-molecule binding using magnetic tweezers. Nucleic Acids Res., 38:7122–7132, 2010. doi: 10.1093/nar/gkq598.

[75] Sergio Cruz-León, Tomáš Majtner, Patrick C Hoffmann, Jan Philipp Kreysing, Sebastian Kehl, Maarten W Tuijtel, Stefan L Schaefer, Katharina Geißler, Martin Beck, Beata Turoňová, et al. High-confidence 3D template matching for cryo-electron tomography. Nat. Commun., 15:3992, 2024.

[76] Tomáš Majtner, Jan Philipp Kreysing, Maarten W Tuijtel, Sergio Cruz-León, Jiasui Liu, Gerhard Hummer, Martin Beck, and Beata Turoňová. cryoTIGER: deep-learning based tilt interpolation generator for enhanced reconstruction in cryo electron tomography. *Commun*. Biol., 8:1443, 2025.

[77] Huabin Zhou, Jan Huertas, M. Julia Maristany, Kieran Russell, June Ho Hwang, Run-Wen Yao, Nirnay Samanta, Joshua Hutchings, Ramya Billur, Momoko Shiozaki, Xiaowei Zhao, Lynda K. Doolittle, Bryan A. Gibson, Andrea Soranno, Margot Riggi, Jorge R. Espinosa, Zhiheng Yu, Elizabeth Villa, Rosana Collepardo-Guevara, and Michael K. Rosen. Multiscale structure of chromatin condensates explains phase separation and material properties. Science, 390:eadv6588, 2025. doi: 10.1126/science.adv6588.

[78] Mikhail V. Bass, Tatiana Nikitina, Davood Norouzi, Victor B. Zhurkin, and Sergei A. Grigoryev. Nucleosome spacing periodically modulates nucleosome chain folding and dna topology in circular nucleosome arrays. Journal of Biological Chemistry, 294(11):4233–4246, 2019. ISSN 0021-9258. doi: 10.1074/jbc.RA118.006412. URL https://www.sciencedirect.com/science/article/piiS0021925820418290.

[79] Kangsun Yun, Jae-Seon So, Arijita Jash, and Sin-Hyeog Im. Lymphoid enhancer binding factor 1 regulates transcription through gene looping. The Journal of Immunology, 183(8):5129–5137, 10 2009. ISSN 0022-1767. doi: 10.4049/jimmunol.0802744. URL https://doi.org/10.4049/jimmunol.0802744.

[80] Jon K. Chen, Claris Y.Y. Chong, and Lu Gan. Toward 3-D genome visualization with *in situ* cryo-electron tomography. Curr. Opin. Struct. Biol., 99:103301, 2026. doi: 10.1016/j.sbi.2026.103301.

[81] Nathan Jentink, Carson Purnell, Brianna Kable, Matthew T. Swulius, and Sergei A. Grigoryev. Cryoelectron tomography reveals the multiplex anatomy of condensed native chromatin and its unfolding by histone citrullination. Mol. Cell, 83:3236–3252.e7, 2023. doi: 10.1016/j.molcel.2023.08.017.

[82] Adrian A. Canutescu and Roland L. Dunbrack, Jr. Cyclic coordinate descent: a robotics algorithm for protein loop closure. Protein Sci., 12:963–972, 2003. doi: 10.1110/ps.0242703.

[83] Yongjia Duan, Ye Lu, Sanjoy Paul, Johannes Betz, Léa P. Wilhelm, Annan S. I. Cook, Xuefeng Ren, Elias Adriaenssens, Sascha Martens, Ian G. Ganley, Gerhard Hummer, and James H. Hurley. Reconstitution of multistep recruitment of ULK1 to membranes in autophagy. Sci. Adv., 12:eaeg3201, 2026. doi: 10.1126/sciadv.aeg3201.

[84] Shuxiang Li, Maria J. Aristizabal, Sergei A. Grigoryev, and Anna R. Panchenko. Molecular dynamics guided all-atom reconstruction of cryo-ET maps reveals mechanisms of histone tail mediated chromatin compaction. Nat. Commun., 17, 2026. doi: 10.1038/s41467-026-77871-9.

