## Supplementary Information for "Gene-scale DNA and chromatin ensembles with atomic resolution"

### Supporting methods

#### Hierarchical Chain Growth (HCG) - Algorithm

HCG-DNA takes as input a target DNA sequence and an atomistic fragment library containing conformational ensembles for all 136 unique double-stranded tetranucleotide sequence contexts. The target sequence is decomposed into overlapping tetranucleotide fragments, shifted by two base pairs along the sequence. Consequently, neighboring fragments share a two-base-pair sequence (Fig. 1 of the main text, and Supporting Fig. 1). For example, the sequence 5′-TTGTAGGCTC-3′ is represented initially by the fragments: 5′-TTGT-3′, 5′-GTAG-3′, 5′-AGGC-3′ and 5′-GCTC-3′ (Supporting Fig. 1).

For each tetranucleotide fragment, a conformation is drawn randomly from the corresponding atomistic library. Neighboring fragments are then assembled pairwise. Because adjacent fragments share two base pairs, the terminal shared region of the fragment closer to the 5′ end is used as the reference for aligning the corresponding region of the fragment closer to the 3′ end. The 5′ fragment remains fixed, whereas the 3′ fragment is translated and rotated by a rigid-body transformation $(\mathbf{R}, \mathbf{t})$ that minimizes the positional RMSD between the heavy atoms in the nucleobase and the ribose carbon atoms of the shared two-base-pair alignment region (Supporting Fig. 1 - level 1).

After alignment, the proposed assembly is accepted only if it satisfies the RMSD criterion and contains no steric clashes. If the RMSD is below 0.6 Å, we check for clashes between adjacent residues using a heavy atom distance cut-off of 2 Å. If no clashes are detected in this step, we then check that the phosphorus (P) atoms in the assembled structure are $> 4$ Å to prevent further clashes. The duplicate copy of the shared sequence is then removed, yielding a single continuous DNA segment. Thus, merging two overlapping 4-bp fragments produces a 6 bp segment. In the example in Supporting Fig. 1, 5′-TTGT-3′ and 5′-GTAG-3′ are merged to produce 5′-TTGTAG-3′, while 5′-AGGC-3′ and 5′-GCTC-3′ are merged independently to produce 5′-AGGCTC-3′.

Assembly then proceeds hierarchically. At the next level, neighboring segments generated at the preceding level are aligned through their shared terminal sequence, checked for steric compatibility, and merged in the same manner. This pairwise procedure is repeated until the full target sequence has been assembled.

To avoid storing the complete atomic coordinates of every intermediate structure, HCG-DNA stores the rigid-body transformations generated at each assembly step. The position of any original fragment in a higher-level segment can be reconstructed by recursively composing the transformations along the hierarchy. Atomic coordinates are generated only when required for alignment, steric-clash evaluation, or writing the final structures.

Independent Monte Carlo workers perform fragment sampling and assembly attempts in parallel. Accepted rigid-body transformations are stored in chunked HDF5 files (“Chunk 1”, “Chunk 2”, etc. Because full-length atomic coordinates are not retained in memory during assembly, the memory requirements remain small even for very long DNA sequences (Supplementary Fig. 4).

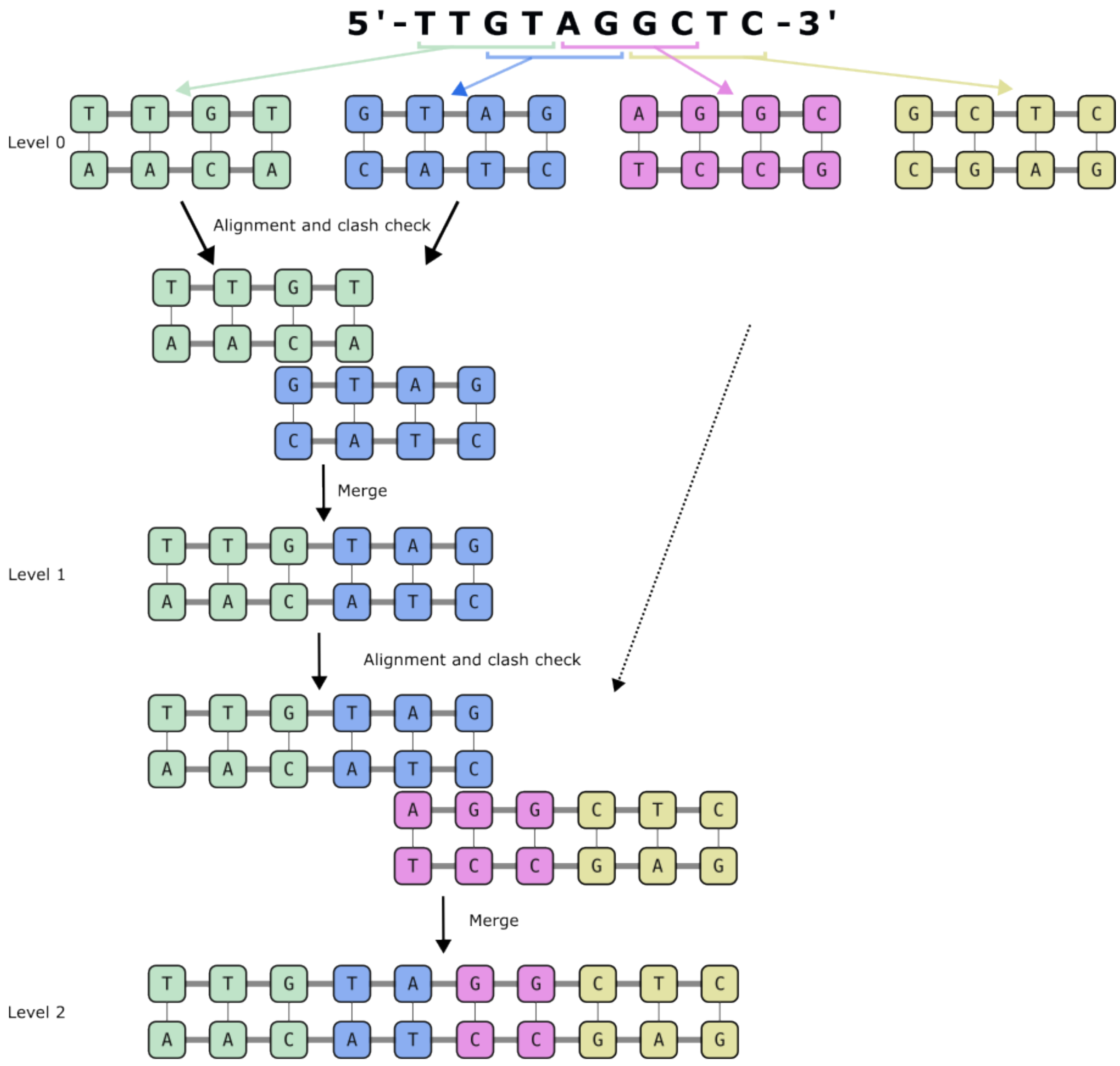


**Supporting Figure 1:** Schematic illustration of the HCG-DNA assembly process for the example sequence 5′-TTGTAGGCTC-3′. First, the target sequence is split into tetranucleotide fragments with an overlap of 2 bp. At each level, fragments are aligned with two overlapping residues and checked for clashes. Accepted pairs inherit the central two base pairs and one additional base pair at each terminus.

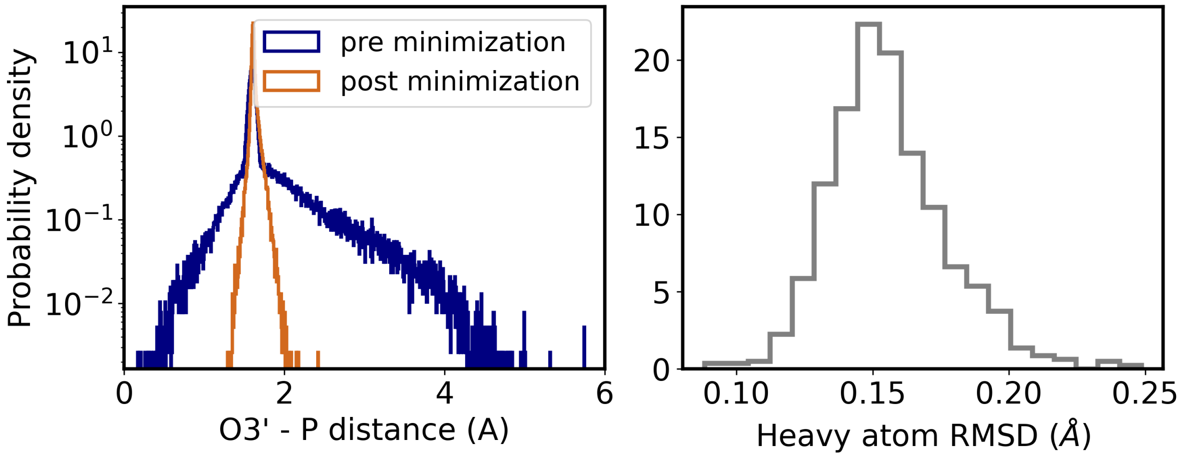


**Supporting Figure 2:** Effects of energy minimization on 1000 HCG structures of 33-bp DNA. **Left** Distribution of O3′-P bonds across the ensemble before (blue) and after minimization (orange). **Right** Heavy atom RMSD across the full chain of individual structures before and after minimization.

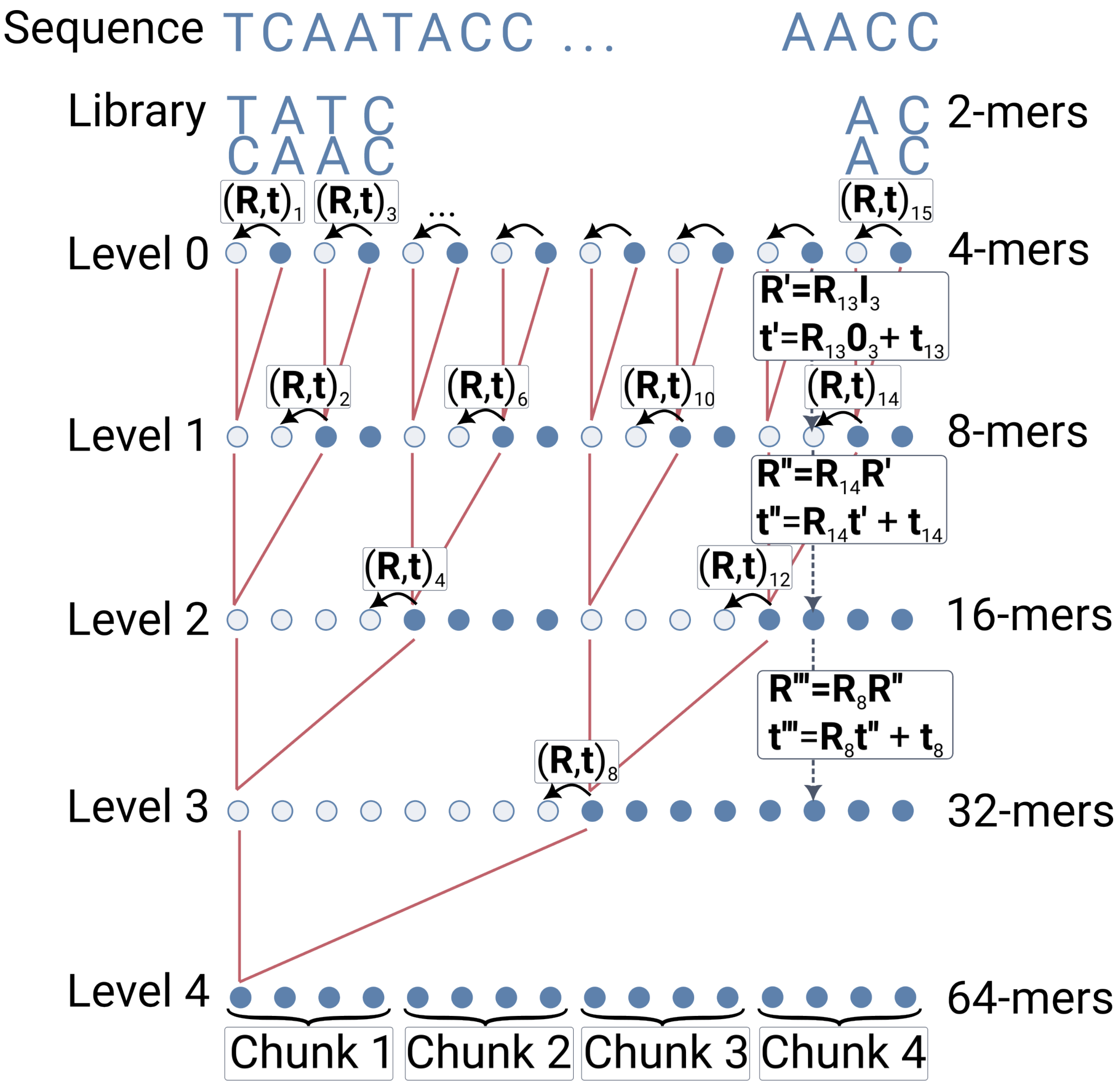


**Supporting Figure 3:** Schematic representation of the HCG algorithm implementation. Starting from a 34 bp sequence that is initially split into 15 2-mer fragments. Each circle represents a 2-mer, where filled circles represent mobile fragments and empty circles represent stationary fragments. The 15 transformations that superpose 2-mer *i* on the adjacent 2-mer *i-1* at any given level are displayed as $(\mathbf{R}, \mathbf{t})_i$, where only the first two and the last transformation on level 0 are shown for clarity. The red lines between levels highlight the $2^n$-mers that are being connected to form a $2^{n+1}$-mers on the subsequent levels.

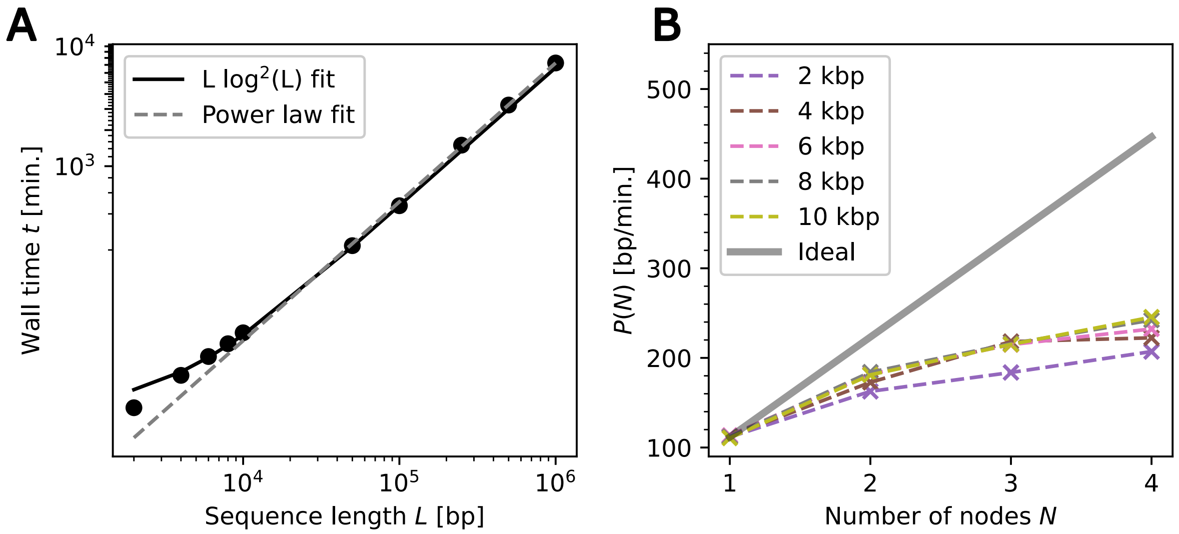


**Supporting Figure 4:** Computational efficiency of the HCG implementation measured in minutes wall time for ensembles of 500 structures. (**A**) Wall time as a function of sequence length $L$ obtained from HCG runs on 4 nodes. Fits were obtained through linear least squares against $L\log^2(L)$ using a first-degree polynomial and through non-linear least squares to a power law $aL^b$. For the fits, only HCG run times for sequence lengths up to 100 kbp (computed on 4 nodes) were used. The optimal parameters for the $L\log^2(L)$ fit are $x_1 = 0.000184(1)$ min./bp and $x_2 = 9.7(14)$ min., RMSE $= 2.5$ min., $R^2 = 0.99$. The optimal parameters for the power law fit are $a = 0.00081(8)$ min./bp and $b = 1.158(7)$. (**B**) Performance $P(N)$ of HCG as a function of number of nodes. The ideal scaling is computed from $P_{id}(N) = P(1)N$ with $P(1)$ being estimated from the 2 kbp sequence HCG run. Numbers in parenthesis are fit errors in the last digits. Note that in typical practical situations, one can take advantage also of trivial parallelization by executing multiple HCG-DNA runs in parallel and initiated with different random seeds.

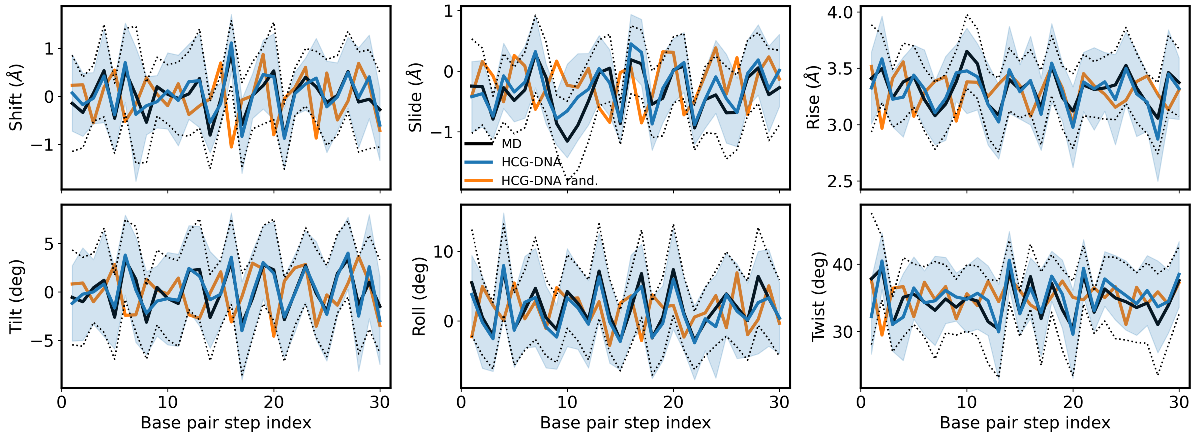


**Supporting Figure 5:** Comparison of helical descriptors between independent MD simulation (black) and HCG ensembles. Position-resolved shift, slide, rise, tilt, roll and twist for a 33-bp DNA duplex from an independent 5-$\mu$s atomistic MD simulation (black) [1] and the corresponding HCG-DNA ensemble (blue). Line and shaded area represent mean and standard deviation across the ensemble, respectively. Terminal base pairs were excluded from the analysis. The orange line shows the ensemble mean obtained from HCG-DNA for an independently generated random 33-bp sequence, illustrating the sequence dependence of the predicted helical parameters. Pearson coefficients of the HCG-DNA of the same sequence (blue) are 0.89 (shift), 0.87 (slide), 0.88 (rise), 0.93 (tilt), 0.94 (roll), 0.80 (twist). For the random sequence (orange) the values are -0.33 (shift), -0.31 (slide), -0.04 (rise), -0.32 (tilt), -0.18 (roll), -0.10 (twist).

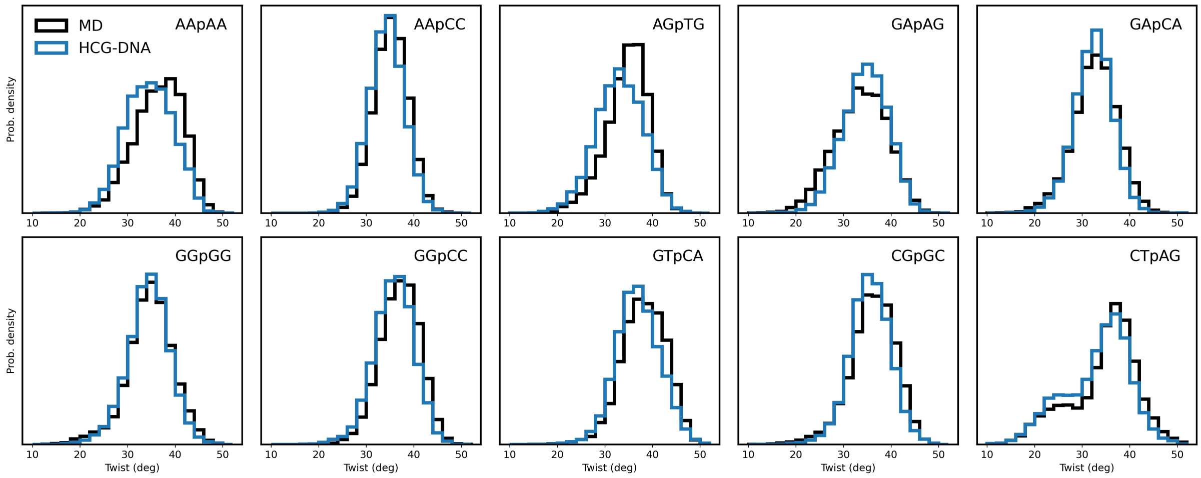


**Supporting Figure 6:** Base pair twist distributions of the central base pair step ($X_2X_3$) of 10 exemplary tetranucleotide sequences embedded in a decanucleotide sequence (5′-GCG$X_1X_2X_3X_4$CGC-3′) from MD simulation and HCG fragment assembly. Sequence of tetranucleotide and central step are indicated.

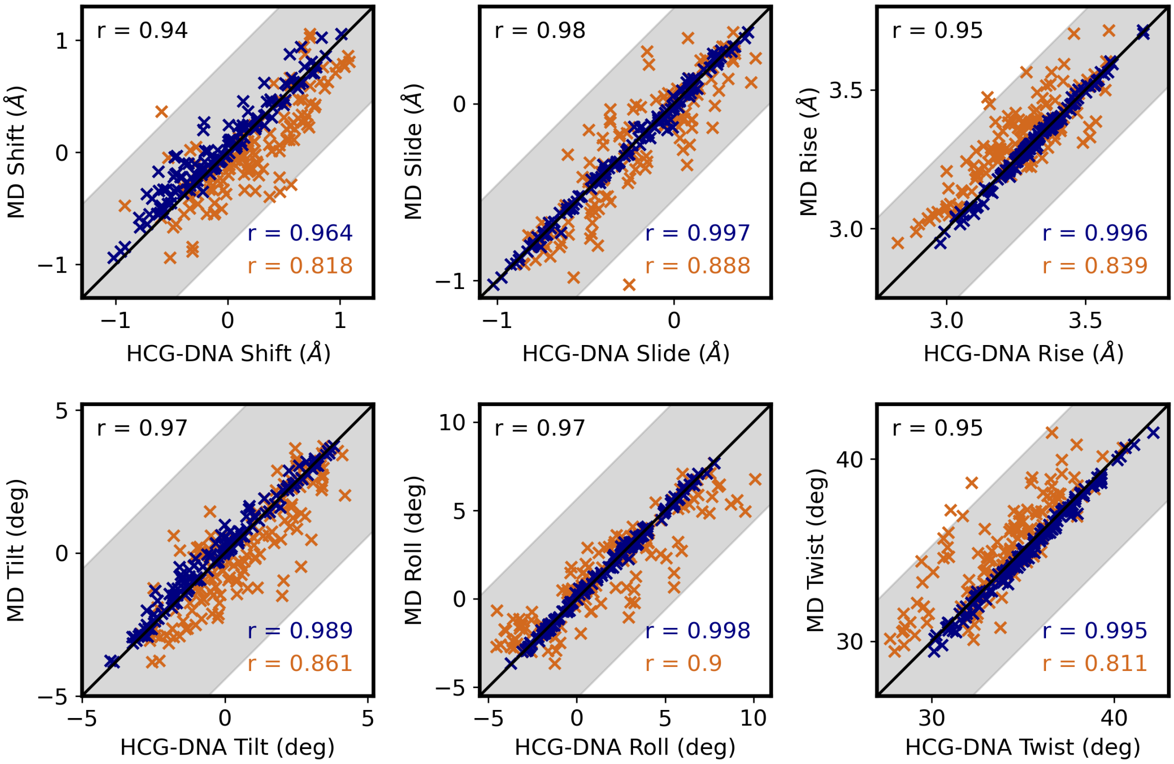


**Supporting Figure 7:** Comparison of base pair parameters shift, slide, rise, tilt, roll and twist of the central base pair step ($X_2X_3$) for all 136 unique tetranucleotides between HCG-DNA ensembles and MD simulation of decanucleotides (5′-GCG$X_1X_2X_3X_4$CGC-3′). HCG-DNA ensembles represented by blue markers were constructed such that the central base pair step is inherited from the fragment simulations by assembling an octanucleotide sequence (5′-CG$X_1X_2X_3X_4$CG-3′). HCG-DNA ensembles represented by orange markers derive the central base pair step from fragment assembly by assembling the same decanucleotide sequence as in MD. Pearson correlation coefficients for each data set are indicated in their respective color. Pearson coefficients of the average of the two datasets (top left) are 0.94 (shift), 0.98 (slide), 0.95 (rise), 0.97 (tilt), 0.97 (roll), 0.95 (twist).

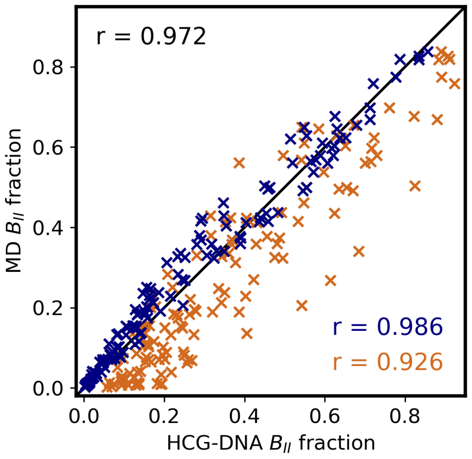


**Supporting Figure 8:** Comparison of fractions of $B_{II}$ fractions at central base pair step of all 136 unique tetranucleotides between HCG-DNA ensembles and MD simulation of decanucleotides (5′-GCGX$_1$X$_2$X$_3$X$_4$CGC-3′). HCG-DNA ensembles represented by blue markers were constructed such that the central base pair step is inherited from the fragment simulations by assembling an octanucleotide sequence (5′-CGX$_1$X$_2$X$_3$X$_4$CG-3′). HCG-DNA ensembles represented by orange markers derive the central base pair step from fragment assembly by assembling the same decanucleotide sequence as in MD. Pearson correlation coefficients are indicated in their respective color. Pearson coefficient of the average of the two datasets is 0.972.

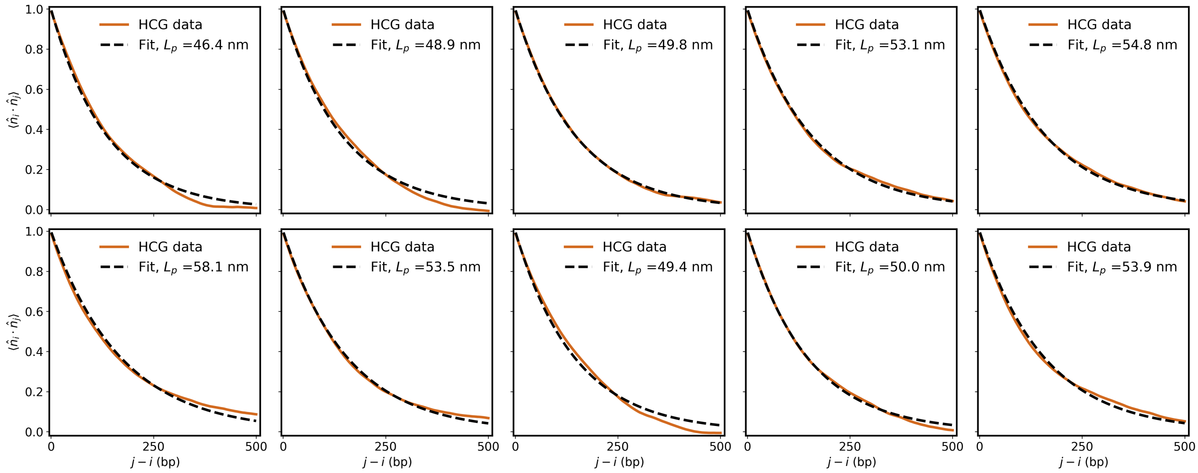


**Supporting Figure 9:** Base pair plane normal vector ($\hat{n}$) correlations between base pairs $i$ and $j$ across separation distances of 10 representative structures from HCG ensemble of full-length $\lambda$-phage genome. Black dashed lines represent exponential decay of the correlations fitted over the full range of 1 to 500. Indicated persistence length in legend correspond to the fit.

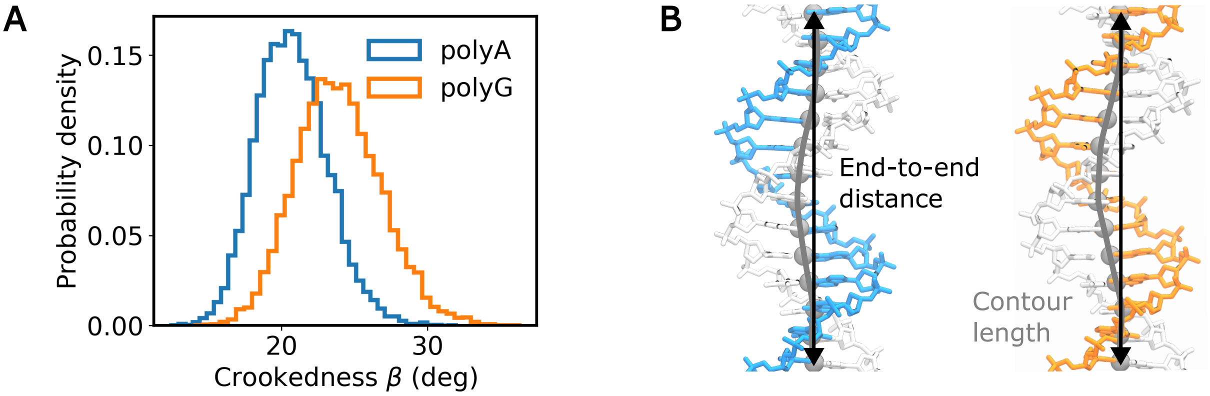


**Supporting Figure 10:** Crookedness analysis of polyA and polyG sequences. (**A**) Distributions from HCG ensembles of 10,000 structures each. (**B**) Average structure of polyA (5′-CGCGAAAAAAAAAAACGCG-3′, left) and polyG (5′-CGCGGGGGGGGGGGCGCG-3′, right) with base pair center of masses indicated by grey spheres with the contour line through them. End-to-end distances are indicated by black arrows.

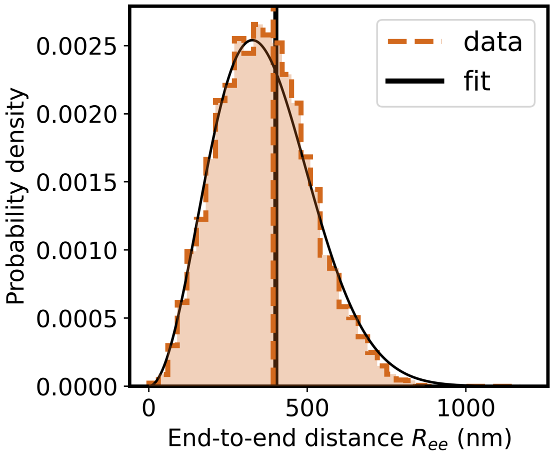


**Supporting Figure 11:** Gaussian chain fit to end-to-end distance distribution of linear pBR322 from HCG-DNA ensembles of 10,000 structures. Vertical dashed line shows moment estimate of $391.7 \pm 1.4$ nm (mean ± s.e.). Black vertical line shows $\sqrt{\langle R^2 \rangle} = 400.3 \pm 2.9$ nm (fit ± s.e.) from gaussian chain fit (main text eq. 1) with error from propagating fit covariance.

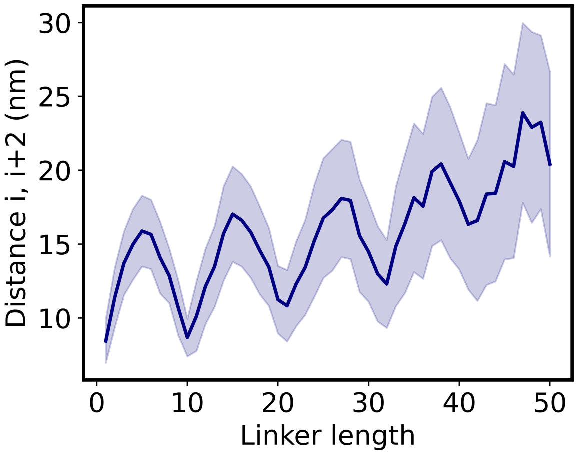


**Supporting Figure 12:** Average distances of nucleosomes i, i+2 in ensembles of 12-nucleosome arrays at linker lengths 1 to 50. Shading represents the standard deviation across 1000 structures.

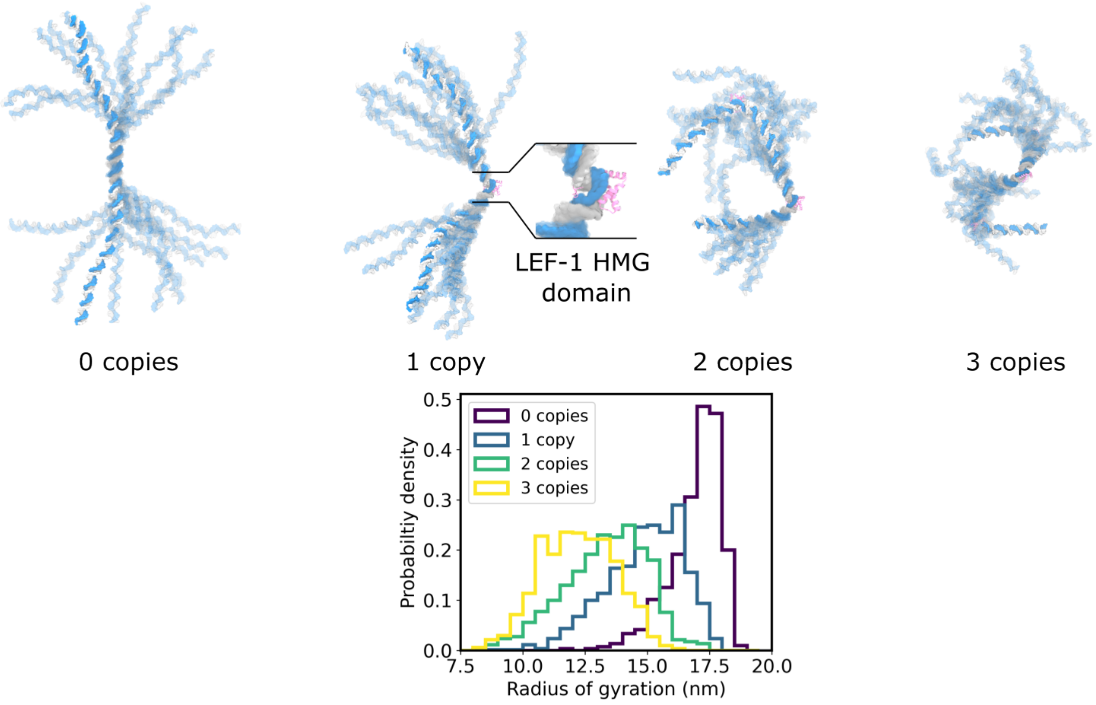


**Supporting Figure 13:** HCG-DNA ensembles of 200 bp duplex with 0 to 3 copies of LEF-1 HMG domain (PDB ID pdb_00002lef[2]) bound.

### References


[1] Sergio Cruz-León, Willem Vanderlinden, Peter Müller, Tobias Forster, Georgina Staudt, Yi-Yun Lin, Jan Lipfert, and Nadine Schwierz. Twisting DNA by salt. *Nucleic Acids Res.*, 50:5726–5738, 2022. doi: 10.1093/nar/gkac445.

[2] John J. Love, Xiang Li, David A. Case, Klaus Giese, Rudolf Grosschedl, and Peter E. Wright. Structural basis for DNA bending by the architectural transcription factor LEF-1. *Nature*, 376:791–795, 1995. doi: 10.1038/376791a0.
